# Convergent gene pairs restrict chromatin looping in *Dictyostelium discoideum,* acting as directional barriers for extrusion

**DOI:** 10.1101/2024.06.12.598618

**Authors:** Irina V. Zhegalova, Sergey V. Ulianov, Aleksandra A. Galitsyna, Ilya A. Pletenev, Olga V. Tsoy, Artem V. Luzhin, Petr A. Vasiluev, Egor S. Bulavko, Dmitry N. Ivankov, Alexey A. Gavrilov, Ekaterina E. Khrameeva, Mikhail S. Gelfand, Sergey V. Razin

## Abstract

*Dictyostelium discoideum* is a unicellular slime mold, developing into a multicellular fruiting body upon starvation. Development is accompanied by large-scale shifts in gene expression program, but underlying features of chromatin spatial organization remain unknown. Here, we report that the Dictyostelium 3D genome is organized into positionally conserved, largely consecutive, non-hierarchical and weakly insulated loops at the onset of multicellular development. The transcription level within the loop interior tends to be higher than in adjacent regions. Loop interiors frequently contain functionally linked genes and genes which coherently change expression level during development. Loop anchors are predominantly positioned by the genes in convergent orientation. Our data suggest that the loop profile may arise from the interplay between transcription and extrusion-driven chromatin folding. In particular, a convergent gene pair serves as a bidirectional extrusion barrier or a “diode” that controls passage of the cohesin extruder by relative transcription level of paired genes.

## Introduction

*Dictyostelium discoideum* (hereinafter referred to as Dicty) is a haploid amoeba which inhabits the forest soil, and undergoes a starvation-triggered developmental program. This program consists of aggregation into a multicellular organism, proto-tissue emergence, and spore formation^1^. Dicty is widely used as a model organism for studying such processes as transcription bursts^2^, chemotaxis^3–5^, cellular signaling^6,7^, autophagy^8^, facultative multicellularity^9^, altruism^10^, and host-pathogen interactions^11,12^.

Dicty’s genome has a low GC-content (22.4%), similar to *Plasmodium*^13^, and up to 10% of its genome consists of repeats. *Dictyostelium intermediate repeat sequences* (DIRS) clusters^14^ seem to serve as centromeres, as they are colocalized with the centromeric histone H3 variant^15^ and feature high levels of H3K9me3^16^. Telomeres are partially formed by rDNA-like elements^17^. Approximately 240 genes are involved in transcription regulation, chromatin remodeling, and histone modifications^18^. Histone marks are generally identical to the ones in animals, with the exception of the H3K27 methylation, which is absent in Dicty^19^. Only a few whole-genome studies of Dicty’s chromatin structure and dynamics in development have been published by now.

A recent work^20^ emphasizes the role of MADS-box transcription factors, like mef2A and srfA, whose deletion causes delays of aggregation initiation and extends the time needed to complete the aggregation. It is worth noting that, the deletion of *smcl1* gene, encoding the cohesin-core subunit, completely blocks the development at the aggregation stage (probably due to impaired mitosis). *Smcl1* gene overexpression also slowed down the formation of multicellular fruit body. Hundreds of long non-coding RNA (lncRNA) loci are potentially involved in multicellular development^21^, and the transcription factor GtaC has been found to play a prevailing role in the process of early aggregation, including cAMP signaling and cell differentiation^22^. Dicty’s chromatin seems to be organized into dinucleosome units with unknown function^23^. However, nothing is known about the 3D organization of the Dicty genome.

The spatial structure of chromatin in most eukaryotes is believed to be predominantly formed by a balance between compartmental segregation and loop extrusion^24^. The former is believed to be a driving force for spatial segregation of active and repressed genome loci, shaped by differences in both physicochemical properties and concentrations of activating and inhibiting chromatin-associated factors^25^. Loop extrusion is an ATP-dependent DNA translocation, mediated by the SMC-family proteins, such as cohesin and condensin. Loop extrusion progresses until the extruder is released or encounters a barrier. One of the thoroughly studied extrusion-limiting factors for cohesin in vertebrates is CTCF (CCCTC-binding factor), which directly interacts with cohesin, thereby stabilizing it on the chromatin^26,27^. At the vast majority of loop anchors in mammalian genomes, CTCF and cohesin co-localize^28^. Other obstacles for loop extrusion are MCM complexes^29^, that have the same cohesin-interacting domain as CTCF^26^, and stalled replication forks^30^. Non-extruding cohesin complexes mediate cohesion of sister chromatids and also appear to be obstacles for the loop extrusion^31^.

Finally, recent studies across various organisms, including bacteria^32^, dinoflagellates^33,34^, fission and budding yeast^30,35,36^, and mammals^37,38^, suggest that transcribed genes can be barriers to loop extrusion. Some studies^39^ suggest that RNA polymerase II itself may act as a semi-permeable barrier for loop extrusion, while others^38,40^ propose that RNA-DNA hybrids (R-loops) could physically interact with cohesin. Although the exact mechanism of this phenomenon remains unknown, it has been noted that highly transcribed genes in convergent orientation are less permeable barriers compared to stand-alone active genes^41^. Additionally, it has been proposed that cohesin might be stalled by macromolecule condensates formed at active promoters and enhancers^42,43^.

Here, we performed *in situ* Hi-C to study the chromatin spatial organization in Dicty during development, coupled with transcriptome profiling, polymer simulations, and molecule dynamics of cohesin which is a putative extruder in Dicty chromatin. Our data suggest that the Dicty 3D genome is organized into loops, which seem to be formed by loop extrusion. Pairs of highly transcribed convergent genes appear to be transcription-tunable barriers for the extrusion. Loops partition the genome into highly transcribed regions, which frequently contain functionally-related genes.

## Results

### Features of the Dicty 3D genome

We analyzed the spatial organization of the Dicty genome using *in situ* Hi-C^28^ in four life cycle stages. These are axenically growing vegetative cells and three stages of starvation-induced multicellular development: migrating cells, early and late aggregates (2, 5 and 8 hours of starvation, respectively; Fig. 1a). To construct Hi-C maps, we used the DpnII restriction enzyme, cutting the Dicty genome into fragments with a median length of 292 bp. Despite a relatively low GC-content of the Dicty genome (22.4%), this length is comparable to the average length of DpnII restriction fragments in organisms with more GC-balanced genomes, such as budding yeast, fruit fly, and human (Extended Fig. 1a). After merging of Hi-C reads from biological replicates (Extended Fig. 1b), we obtained 46.6-55.2 million unique contacts (Supplementary Table 1) that enabled us to analyze the Dicty 3D genome with the resolution up to 500 bp.

**Figure 1.**
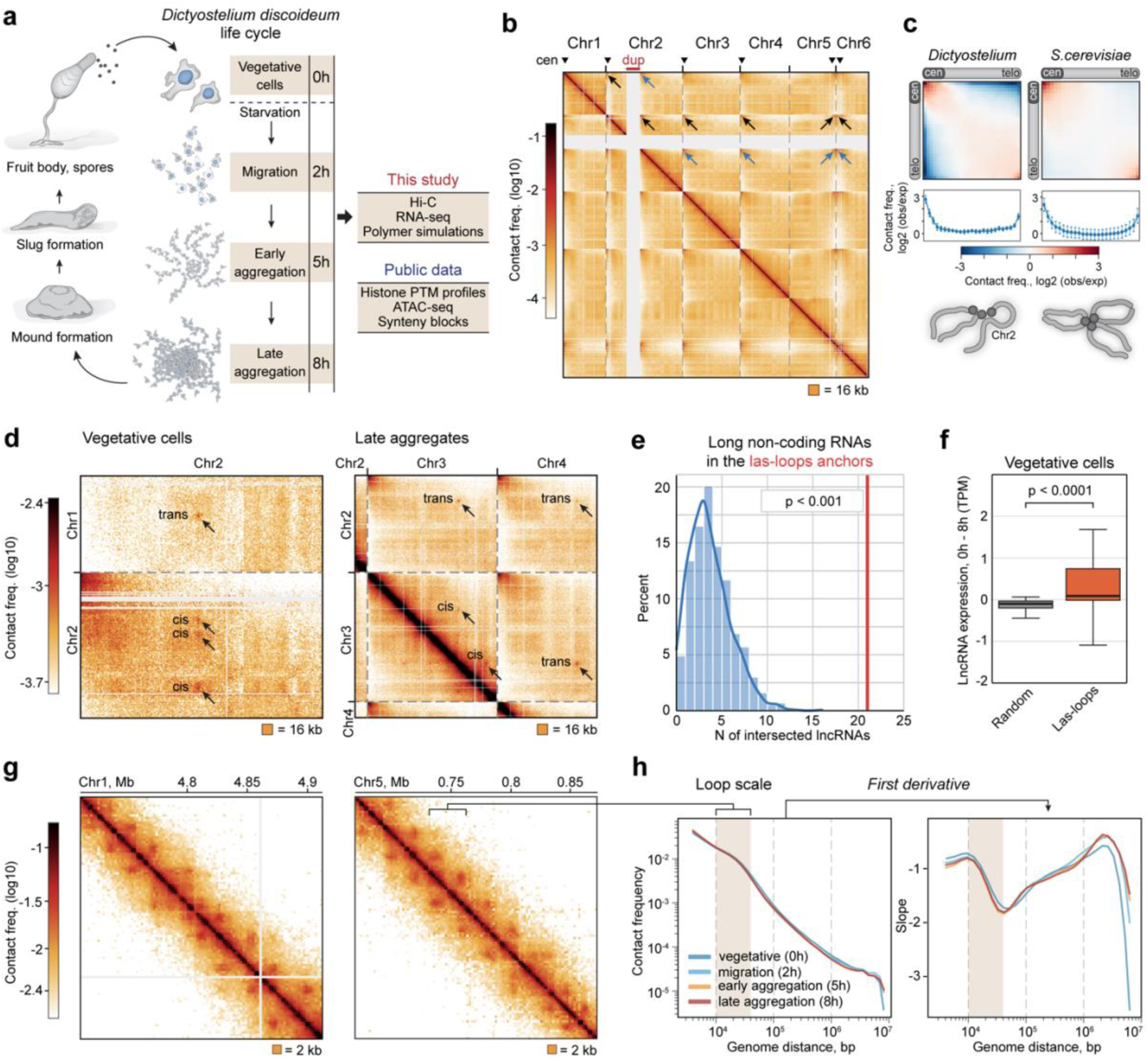
Features of Dicty 3D genome. **a**, Short scheme of the Dicty multicellular development. **b**, Whole-genome Hi-C map. Centromeres are marked with triangles. Arrows show contacts between centromeres and the DIRS-containing region downstream of large duplication on chr2 (marked with red). Note that *Dictyostelium* chromosomes are acrocentric. **c**, Rabl configuration averaged over all chromosomes in Dicty and *S. cerevisiae*. The profile of contact frequency along the main diagonal is shown below the heatmap. **d**, Representative examples of large-scale loops (las-loops). The resolution is 16 kb. **e**, Enrichment of lncRNAs in las-loop anchors. *p*-value in the permutation test. **f**, Difference of the expression level between vegetative cells (0h) and late aggregation stage (8h) for randomly picked lncRNAs (gray) and lncRNAs from los-loop anchors identified in vegetative cells. **** *p-*value in the MWU. **g**, Contact maps demonstrating the presence of regular loops. The resolution is 2 kb. **h**, *P_c_*(*s*) (left panel) and its first derivative (right panel) for the development stages. The range of genomic distances corresponding to loop sizes is indicated as “loop scale”.

Whole-genome maps show that Dicty chromosomes are weakly segregated from each other and establish prominent *trans* contacts (Fig. 1b). The strongest contacts are formed between chromosome termini enriched with DIRS, whose clusters likely serve as centromeres (Fig. 1b, black arrows)^17^. The AX4 strain carries a large inverted duplication on chr2 (Fig. 1b, highlighted in red). The 3’-end of this duplication contains a partial copy of the DIRS element^17,44^, and this region interacts with the DIRS-containing termini of other chromosomes (Fig. 1b, blue arrows). Thus, Dicty chromosomes are arranged in an unusual Rabl configuration, where chr2 forms a large loop between DIRS-containing loci (Fig. 1c). This configuration is preserved during all analyzed development stages (Extended Fig. 1c).

As revealed by both visual inspection and compartment analysis using principal components, Dicty chromosomes lack a detectable partitioning into A/B compartments^45^ (Extended Fig. 1d), similar to yeast chromosomes. This is likely caused by the high density of TSSs: 72, 51, 11, and 1 TSSs per 100 kb on average for Dicty, budding yeast, fruit fly, and human cells, respectively. As a consequence, Dicty genome is not apparently “barcoded” with large active and repressed regions.

However, we observed numerous spot contacts which are formed between loci separated by several Mb and even *in trans* (Fig. 1d). Using manual annotation, we identified 12 and 11 <u>la</u>rge-<u>s</u>cale contacts (further referred to as *las-loops*) in vegetative cells and late aggregates, respectively (Supplementary Table 2). Notably, those las-loops are visually absent in migrating cells and early aggregates. A remarkable feature of the las-loops is the formation of a rosette-like structure, where several loci interact with each other *in cis* and *in trans* (up to 9 las-loops in the rosette). All identified las-loops are stage-specific (Extended Fig. 1e), this implies that the las-loop profile could be linked to regulation of cell physiology. Indeed, we found that the las-loop anchor loci are significantly enriched (*p* < 0.001, Mann-Whitney U test, MWU) with genes of long non-coding RNAs (lncRNAs), previously identified in Dicty^21^ (Fig. 1e). At a specific development stage, only a minor fraction of lncRNA loci are involved in the las-loop network, but the majority of las-loop anchors (71%) contain at least one lncRNA gene (Supplementary Table 2). Moreover, the las-loop formation correlates with an active transcription state of lncRNA genes located in the anchor loci (Fig. 1f). This suggests that the transcription of at least some lncRNA genes in Dicty cells is controlled by the formation of active chromatin hubs^46^ or transcription factories^47^.

### Arrays of consecutive loops as a distinctive feature of the Dicty 3D genome

At short genomic distances, Dicty chromatin is routinely organized into short loops that can be easily seen by visual inspection of the heatmaps (Fig. 1g). Accordingly, the analysis of the contact probability dependence on the genomic distance *P_c_*(*s*) has shown a hump at the distance of 10-40 kb for all development stages (Fig. 1h). To systematically analyze the loop profile, we annotated loops by both LASCA^48^ and Chromosight^49^. The intersection of their outputs provided an annotation that aligned closely with visual assessments. For each development stage, we identified ∼1,300 loops, which covered 65-68% of the genome (Supplementary Table 3). These loops display unique properties that differentiate them from typical patterns of chromatin folding in other organisms. To contrast the features of loops in Dicty, we compared them to cohesin-dependent interphase loops in human^50^, condensin-dependent S-phase loops in yeast^51^, and potentially unrelated to loop extrusion *topologically associated domains* (TADs) in Drosophila^52^.

Firstly, the loop profile in Dicty is nearly identical at all development stages (Fig. 2a), with the majority of discrepancies between stages caused by false positive and false negative calls. Secondly, the loop size varies in a surprisingly narrow range from 10 to 40 kb (Fig. 2b, left panel; Extended Fig. 2a). This sharply contrasts with the data from human and fruit fly cells, where the loops and TADs identified at a comparable map resolution vary from 40 kb to 3 Mb (75-fold^28^) and from 3 kb to 460 kb (153-fold^53^), respectively. Finally, about 70% of Dicty loops are organized into sequential arrays, with up to 10 loops per array (Fig. 2b, right panel).

**Figure 2.**
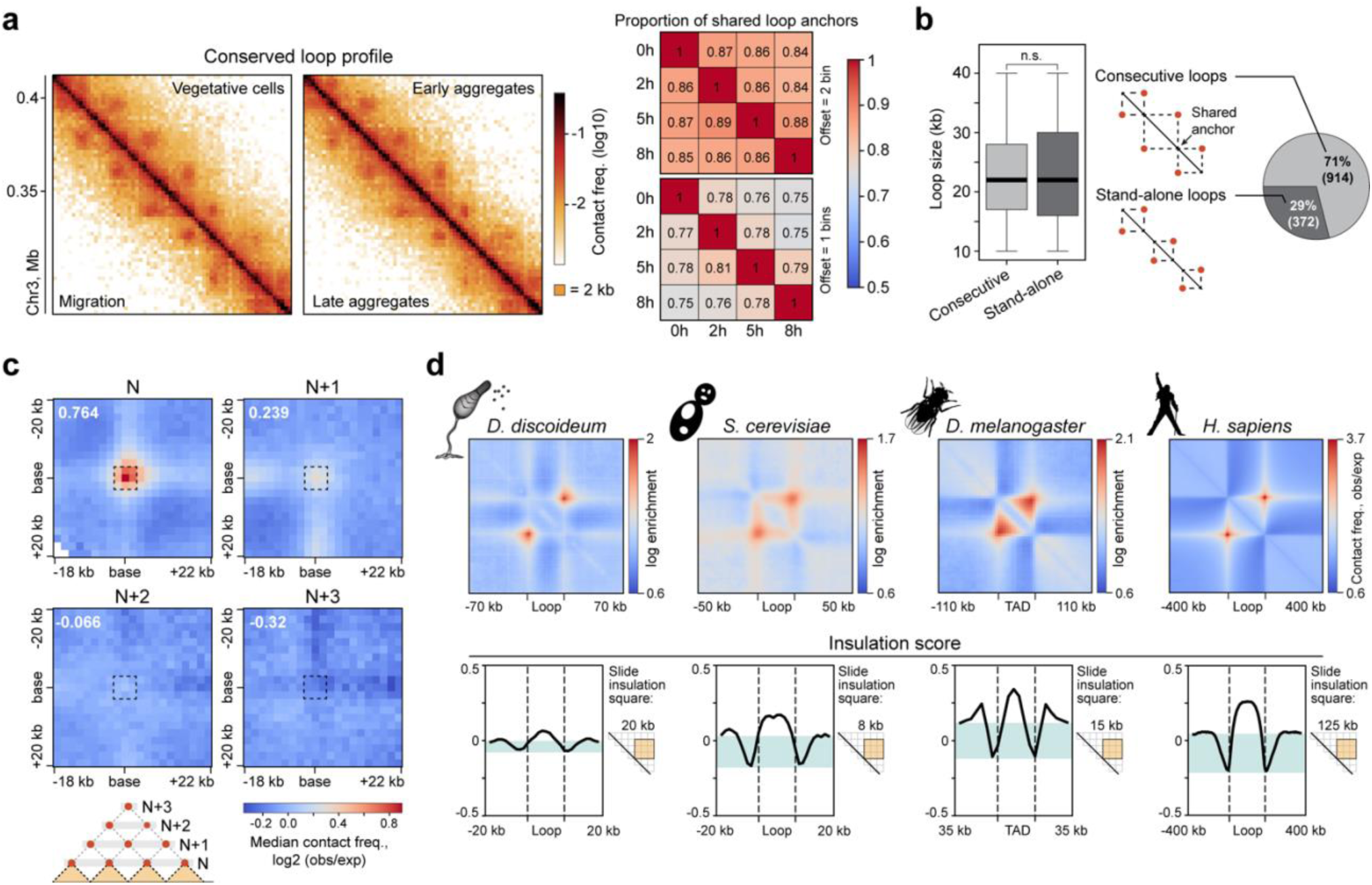
Dicty chromosomes are partitioned into largely consecutive, non-hierarchical and weakly insulated loops. **a**, Left: representative example of a region with conserved loop profile. The resolution is 2 kb. Right: proportion of loop anchors conserved in pairwise comparisons of development stages. **b**, Proportion and size distribution of consecutive and stand-alone loops in vegetative cells, n.s. - non-significant difference in the MWU. **c**, An averaged loop between adjacent anchors (N), and between anchors separated with one (N+1), two (N+2), or three (N+3) anchors. **d**, Averaged loops / TADs (upper panel) and median profile of the insulation score (IS) in different organisms. The dashed lines show loop anchors / TAD boundaries. The size of the sliding window for the IS calculation is shown to the right of the profiles. Data source: *S. cerevisiae* - Micro-C XL in S-phase cells^51^; *D. melanogaster* - Hi-C in spermatogonia^52^; *H. sapiens* - Hi-C in GM12878 lymphoblastoid cells^50^.

Within these arrays, adjacent loops have a shared anchor and hereinafter are referred to as *consecutive loops*. A remarkable feature of consecutive loops is an almost complete absence of hierarchical interaction between their anchors. In other words, each anchor within an array forms a loop with only the two nearest partners (Fig. 2c). This suggests that only a subset of loops is formed in each individual cell, and that the presence of adjacent loops in the same cell is infrequent. In contrast, loops in human^54^ and TADs in fruit fly^55^ interphase cells, as well as loops in budding yeast S-phase cells^56^ are often nested: the large TADs and loops are composed of smaller ones. Finally, unlike the TAD boundaries and anchors of yeast S-phase loops, Dicty’s loop anchors are weak insulators (Fig. 2d and Extended Fig. 2b). This is manifested in the enrichment of contacts inside human and yeast loops, and fruit fly TADs, but not inside Dicty loops (Fig. 2d). Thus, Dicty loops are not compacted into self-interacting domains.

### Functionally linked genes grouped within loop interiors

We proceeded with the analysis of biological relevance of chromatin looping for the Dicty genome functioning. As revealed by previous studies^22,57,58^ and our RNA-seq data (Extended Fig. 3a, b, Supplementary Table 4), even the immediate early stages of the Dicty multicellular development, such as starvation and aggregation, are accompanied by large-scale changes in gene expression program. To characterize these changes, we described expression of each gene during development as a series of transitions between consecutive stages, or *development trajectory*^57^. We performed k-means clustering of gene expression values and identified four robust clusters of genes with similar development trajectories (Fig. 3a), hereinafter named *trajectory gene clusters* (TGCs).

**Figure 3.**
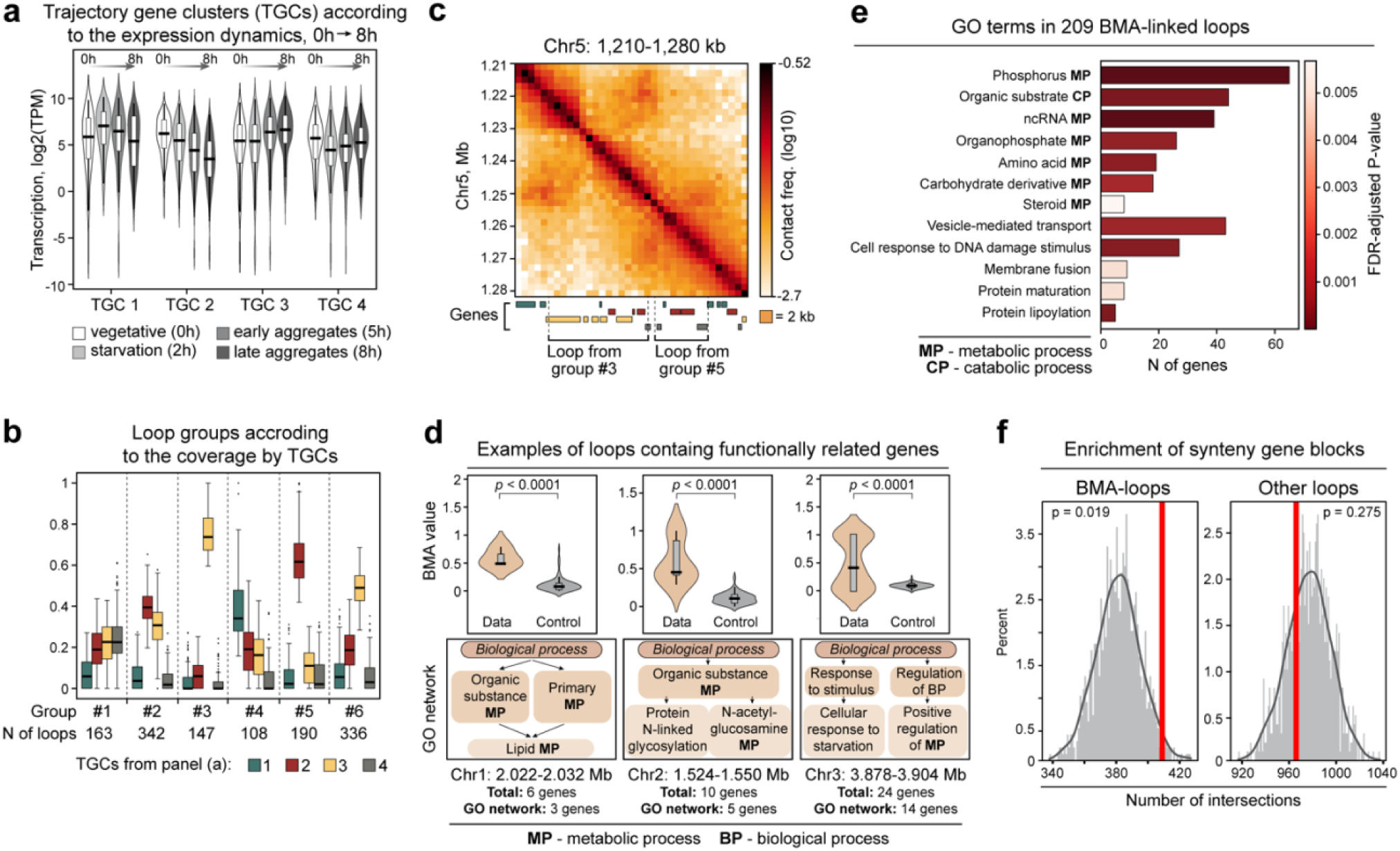
Loops are functional domains of the Dicty genome. **a**, Trajectory gene clusters (TGCs) identified according to changes in expression level during development (development trajectories). **b**, Coverage of genes from different TGCs within loop groups identified by k-means, with coverage by TGC as an input. The number of loops in each cluster is indicated. **c**, Representative examples of loops preferentially covered by genes from different TGCs (gene colors correspond to TGC colors from panel **b**). **d**, Representative examples of loops containing functionally related genes as revealed by the best match averaging (BMA) analysis of GO terms. The higher is the BMA value of a loop, the more prominent are functional relationships between genes within the loop. *P*-value is calculated with the permutation test. The key terms from the GO network of the loop are shown. **e**, GO terms of 167 loops containing functionally related genes, according to the BMA analysis. **f**, Enrichment of synteny gene blocks (*D. discoideum vs D. purpureum*) in BMA-loops and all other loops. Synteny block coordinates are taken from^59^. Permutation test p-values are shown.

Loops in vegetative cells could be clearly grouped according to the coverage by genes of these four TGCs (Extended Fig. 3c; a principal scheme of genes clustering into TGCs and grouping of loops based on coverage by different TGCs is shown in Extended Fig. 3d). Loop groups #2, #3 and #5 are of particular interest since they harbor almost exclusively the genes whose expression tends to either gradually decrease (genes belonging to TGC-2, red boxplots) or increase (genes belonging to TGC-3, yellow boxplots) upon the progression from vegetative cells to late aggregates (Fig. 3b, c).

Further, among the loops identified in vegetative cells, 167 loops (13%) contained genes with highly similar networks of gene ontology (GO) terms, as revealed by the best match average (BMA) analysis (BMA-loops; Fig. 3d, Supplementary Table 5; see Methods for the details). These GO networks are enriched with terms related to phosphorus, amino acids, carbohydrates, and steroid metabolic processes, as well as to vesicle-mediated transport, and response to DNA damage (Fig. 3e), suggesting the potential role of these genes in the response to changes in environmental conditions. Interestingly, BMA-loops are enriched with synteny gene blocks previously identified in the comparison of *D. discoideum* and *D. purpureum* genomes^59^ (Fig. 3f). Together, these observations indicate the presence of functionally related and evolutionary conserved gene islands within the Dicty genome, and that such islands are demarcated with loop anchors.

### Convergent gene orientation at loop anchors

We analyzed the functional properties of Dicty loops. The preliminary visual inspection of the Hi-C maps revealed different types of loops. Some of them have a nearly symmetrical circular shape, while others are clearly elongated along one of the Hi-C map axes (see Fig. 1g and Fig. 2a for examples). To identify such “<u>el</u>ongated loops” (referred to as el-loops) systematically, for each loop, we calculated the ratio between the sums of contacts along the two Hi-C map axes (Fig. 4a, b, see Methods for the details) and separated loops into three categories (Fig. 4c, left panel): symmetrical (ordinary loops), elongated in the 3’-direction and elongated in the 5’-direction.

**Figure 4.**
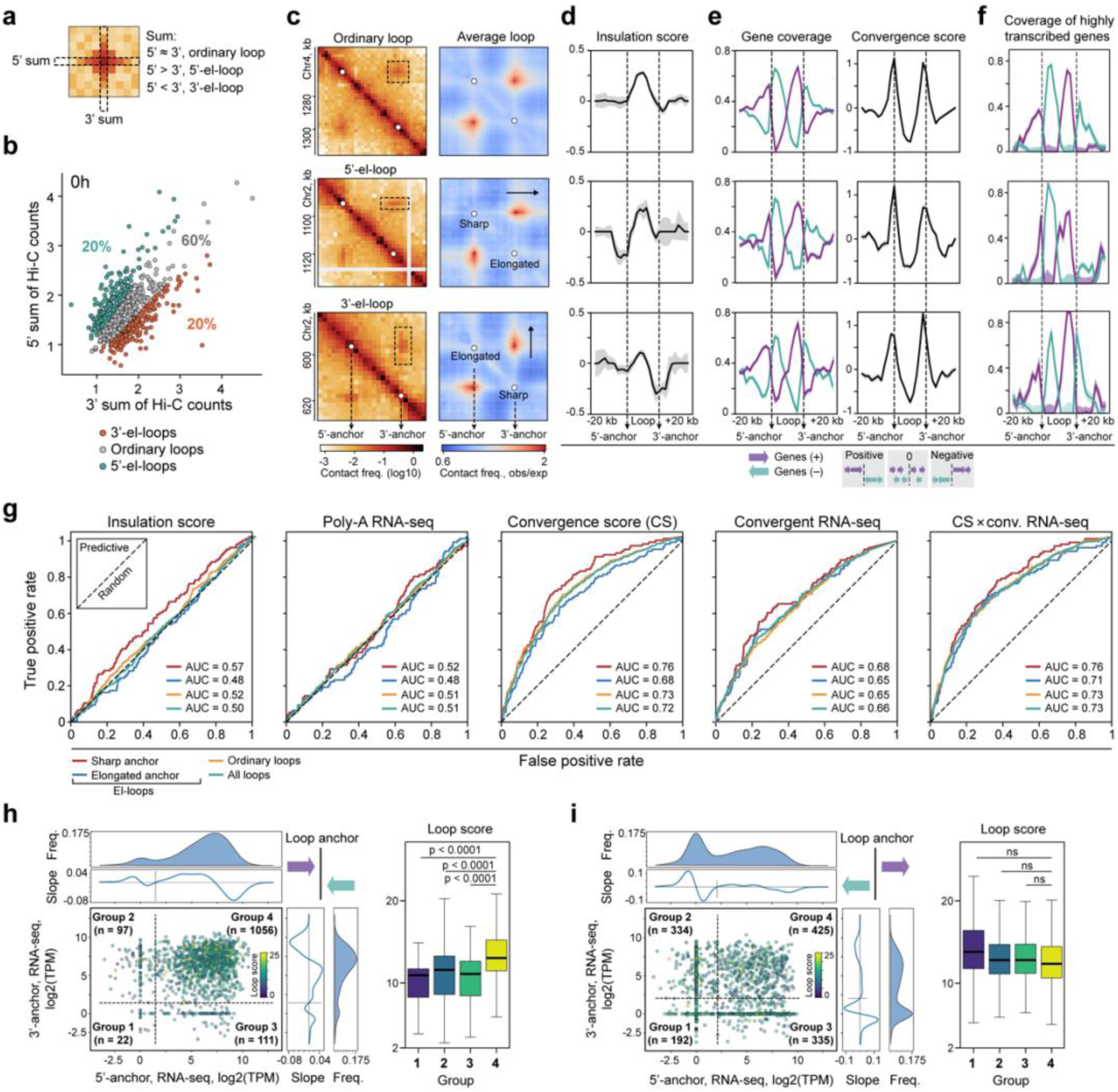
The loop anchor positions are determined by convergently oriented genes. **a**, Scheme of the Hi-C count calculation for the identification of elongated loops (el-loops). **b**, Distribution of sums of Hi-C counts for individual loops in vegetating cells. **c**, Left: representative examples of ordinary loops and el-loops. The resolution is 2 kb. Right: averaged ordinary loops and el-loops. Sharp and elongated anchors of el-loops are indicated **d**, Median profiles of the insulation score in vegetating cells. Bootstrap standard error is indicated. **e**, Median profiles of the gene coverage (left) and convergence score (right). Genes transcribed from the plus and minus strand are shown in violet and aqua, respectively. **f**, Median coverage of genes with high transcription level (above the median in vegetating cells). **g**, Prediction of loop anchors of different types in vegetating cells using logistic regression models based on the genome-wide profiles of IS, transcription (Poly-A RNA-seq), convergence score (CS), transcription of convergent genes (Convergent RNA-seq), and RNA-seq-weighted CS. The receiver operating characteristics (ROC curves) and the AUC (area under the curve) values are shown. **h,** Left: scatterplot showing the level of convergent transcription in a 2-kb vicinity of loop anchors. Dots are colored according to the loop score calculated as a sum of the obs/exp Hi-C counts in the 3×3 pixels area around the pixel identified as a loop (the higher is the score, the brighter is the loop in the Hi-C map). The thresholds of the loop score (dashed lines) are selected as arguments of the minima of the density profiles (Freq. curves shown at the top and to the right of the scatter plots). n is the number of loops in the group. Right: Loop score distribution within loop groups defined in the left panel. *p*-values in the MWU. **i,** The same as (**h**), but based on the level of divergent transcription. n.s. - non-significant difference in the MWU.

To check robustness of el-loop calling, we averaged subsections of Hi-C map over el-loop coordinates (Fig. 4c, right panel). This resulted in strong insulation at the sharp anchors of el-loops, while elongated anchors of el-loops and anchors of symmetrical loops do not insulate well (Fig. 4d; Extended Fig. 4a). Nothing is known about the presence of the functional homologs of mammalian and *Drosophila* insulator proteins in Dicty. We hypothesized that insulation barriers could be established by moving polymerase on DNA, as has been previously shown in bacteria^32^, yeast^36^, and some higher eukaryotes^38^. We thus tested the possibility that the loop morphology (as well as loop formation *per se*) could be linked to the transcription at the anchor loci.

Firstly, we found that the gene order is remarkably non-random in both el-loops and ordinary loops. Inside loops, genes transcribed from the plus strand are primarily located at the 3’-anchor, while genes transcribed from the minus strand are overrepresented at the 5’-anchor, and outside a loop the arrangement of genes is opposite (Fig. 4e, left panel; Extended Fig. 4b). To estimate general tendency in gene orientation, we calculated convergence score^60^ and observed a strong peak of convergence score at the median profile around loop anchors (Fig. 4e, right panel; Extended Fig. 4c). Therefore, the loop anchors are sites of convergent gene orientation, which is reminiscent of recent findings in dinoflagellates where the contact domain boundaries coincide with the junctions between convergent gene arrays^33,34^.

Secondly, we noted that looped architecture is linked to the gene activity at the convergent positions. Genes adjacent to the loop anchors are actively transcribed, and the transcription level is higher at sharp insulating anchors of el-loops (Fig. 4f; Extended Fig. 4d). Notably, this pattern is not restricted to the loops only, since the insulation score at a given genomic bin significantly anti-correlates with the transcription level within this bin (Spearman’s *ρ* = −0.47, *p* < 0.01; Extended Fig. 4e; we should note the large negative values of insulation score mean low number of contacts between flanks of a locus, and thus means high insulation at a locus). In particular, this correlation is prominent at TSSs of differentially expressed genes, which hints at a possible role of RNA-polymerase II as a barrier for spatial interactions in chromatin (Extended Fig. 4f).

As a whole, these observations suggest that convergently oriented genes and high transcription levels could be contributing to the determination of the loop anchor positions. To test this hypothesis, we applied logistic regression based on the combination of insulation level, gene convergence, and transcription level to predict loop anchor positions (Fig. 4g). Unexpectedly, both the insulation score and the transcription level were poor predictors for positioning of anchors, including insulating anchors of el-loops. This suggests that high transcription and insulation at the genomic bin are insufficient to create a prominent chromatin loop. Although the transcription level of convergent pairs is a slightly better predictor than the total transcription level, the best predictor for all types of anchors is the convergence score *per se* (Fig. 4g, middle panel). Interestingly, the combination of the transcription level of convergent genes and the convergence score produces only a mild improvement of predictions (Fig. 4g, rightmost panel). Finally, we noticed that the high level of convergent (Fig. 4h, Extended Fig. 4g), but not divergent transcription (Fig. 4i) is associated with high loop strength.

### Convergent gene pairs as highly transcribed genome units

Since loop anchors are sites of convergent gene orientation, and convergent transcription is associated with high loop strength, we further focused on the detailed analysis of *convergent gene pairs* (CGPs) in the Dicty genome. In total, we identified 4,672 CGPs (Supplementary Table 6; see Methods for the details) that are shown to be significantly enriched at loop anchors (*p* < 0.001, permutation test) and underrepresented in the loop interiors (Extended Fig. 5a). On average, CGPs are transcribed 2.2-fold higher than other genes (*p* < 0.0001, MWU; Fig. 5a). *Anchor-associated* CGPs that are located in the 4-kb vicinity of any loop anchor demonstrate 12-fold higher transcription (*p* < 0.0001, MWU). Inside loops, CGPs and other genes are transcribed at a similarly high level, while outside loops both are less active, but again, transcription in CGPs is 4.4-fold higher than in other genes (*p* < 0.0001, MWU). Finally, when comparing genes that are not organized in CGPs (non-convergent genes), their average transcription level is ∼10-fold higher inside loops as compared to anchors (*p* < 0.0001, MWU) and loci outside of loops (*p* < 0.0001, MWU; beige violin plots in Fig. 5a). This suggests that (i) CGPs in the Dicty genome form gene complexes which ensure relatively high transcription levels regardless of the genome location, and (ii) Dicty loops are chromosome neighborhoods, which promote transcription of non-convergent genes.

**Figure 5.**
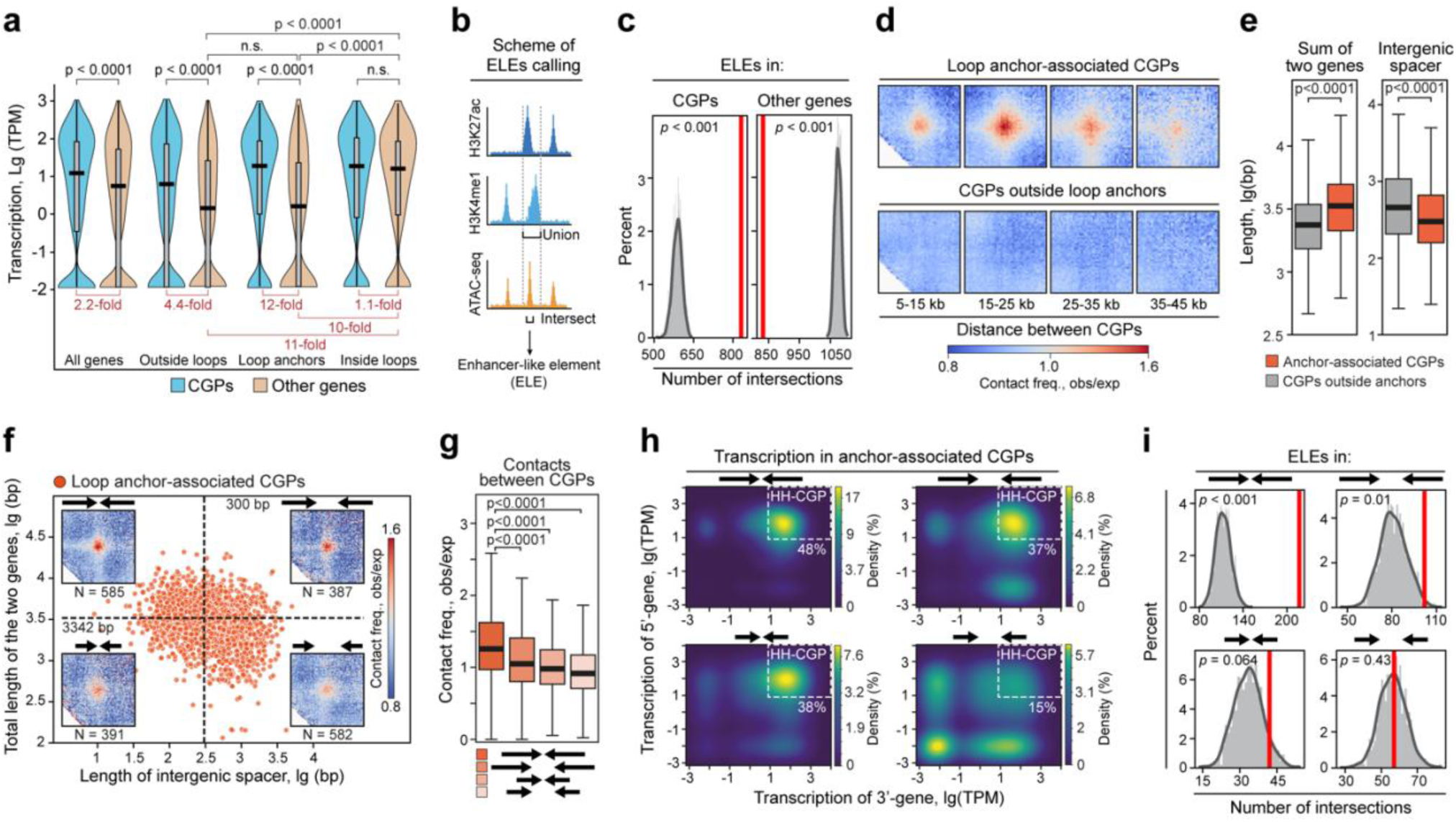
Structural and functional properties of convergent gene pairs in vegetative cells. a, Transcription level of CGPs and other genes located in different positions along the genome relative to loops. *p*-value in the MWU; n.s. - non-significant difference. The fold changes between the median transcription level of convergent gene pairs and other genes are shown in red. **b,** Scheme of the enhancer-like element (ELE) annotation by intersection of ATAC-seq peaks with the union of H3K27ac and H3K4me1 peaks. Reanalyzed data from^20^. **c,** Enrichment of ELEs in CGPs and all other genes. *p*-value in the permutation test. **d,** Averaged interaction between CGPs separated by different genomic distances. **e,** Total length of the two genes (left panel) and length of intergenic spacers (right panel). *p*-values in the MWU. **f,** Distribution of the intergenic spacer length and the total length of two genes in anchor-associated CGPs. The median values are indicated and shown by dotted lines. Averaged contacts between CGPs are shown at the corners. Schematic representation of CGP composition and the numbers of CGPs and are shown above and below the heatmaps, respectively. **g-i,** Pairwise contact frequencies (panel g), kernel density estimate (KDE) plots of transcription level (panel h) and enrichment of ELEs (panel i) in four groups of CGPs assigned in panel (f). The white dotted rectangles in panel (h) demarcate CGPs where TPM > 8 for both genes (HH-CGPs). *p*-values in panels (g) and (i) are calculated in the MWU and permutation test, respectively.

We assumed that high transcription levels in CGPs could be determined by the presence of regulatory elements similar to enhancers in higher eukaryotes. To our knowledge, enhancers in Dicty have not yet been systematically identified, although described for certain genes^61–64^. To create a list of putative *enhancer-like elements* (ELEs), we reanalyzed previously published^20^ ChIP-seq data for H3K27ac and H3K4me1, and chromatin accessibility measured by ATAC-seq (a scheme of the ELE calling is shown in Fig. 5b), epigenetic features of enhancers in higher eukaryotes^65^ frequently used for enhancer *de novo* identification, for example, by ENCODE consortium^66^. In total, vegetative cells have 1,165 putative ELEs with the median length of 984 bp (Extended Fig. 5b, c, Supplementary Table 7). ELE-containing genes are transcribed at a significantly higher level compared to other genes (11-fold, on average; *p* < 0.0001, MWU, Extended Fig. 5d), suggesting that ELEs are positive regulatory elements. ELEs are overrepresented in CGPs and depleted in non-convergent genes (Fig. 5c). This probably explains the high transcription level in CGPs.

We have noticed that many highly transcribed CGPs are located outside of loop anchors and that contact frequency between these CGPs does not exceed the expected values across a wide range of genomic distances (Fig. 5d), resembling divergent gene pairs (DGPs, Extended Fig. 5e). We searched for specific characteristics distinguishing anchor-associated CGPs, and found two key properties of anchor-associated CGPs: significantly shorter intergenic spacers (1.5-fold, *p* < 0.0001, MWU) and longer genes (1.4-fold, *p* < 0.0001, MWU; Fig. 5e).

Among anchor-associated CGPs, the long-gene and short-spacer CGPs form stronger interactions (Fig. 5f, g), have a higher expression (Fig. 5h), and are enriched with ELEs (Fig. 5i). Thus, we concluded that ELE-containing convergent pairs of long, highly transcribed genes with short intergenic spacers are fair determinants of loop anchors.

### Convergent gene pairs shaping chromatin looping

We next asked what mechanism could potentially explain the formation and properties of loops observed in Dicty, including (i) the presence of scaling hump in Hi-C data, (ii) relatively narrow range of loop size, (iii) lack of notable loop hierarchy, (iv) the presence of highly transcribed CGPs of long genes with short spacers in loop anchors. Loop extrusion is a well-studied mechanism that is likely to be present in Dicty as well as in a wide range of species, from bacteria to mammals^32,67^. In mammals and yeast, extrusion establishes a regular loop pattern along the genome^28,51^. This mechanism involves an extruder complex that loads onto DNA and creates a progressively increasing loop connecting two genomic regions until the extruder detaches from DNA and starts the next cycle of extrusion in another location^68,69^. Moreover, transcribing polymerase can interact with the extruder^32,38^, leading to the redistribution of extruders in the direction of moving polymerase^30^.

We hypothesized that convergent transcription at CGPs creates a bidirectional trap for the moving extruder, which can explain the formation of loops between the neighboring CGPs. Typical extruders in higher eukaryotes are cohesin and condensin. Key subunits of the cohesin complex are present in Dicty and are expressed at comparable high levels in all studied development stages (Supplementary Table 8). Previous studies have observed the accumulation of cohesin at the convergent genes in yeast^35,70^, and looping between convergent genes in CTCF-WAPL DKO (double-knockout) cells of mammals^38^. To study whether transcription at CGPs can shape the chromatin looping, we first focused on the subset of CGPs with both genes having high expression (‘high-high CGP’, HH-CGP, TPM > 8 for both genes). On averaged Hi-C maps, HH-CGPs form contact domains with boundaries at the gene promoters (Fig. 6a) and establish stripes of interactions^68,71^ in both upstream and downstream directions (Fig. 6b). This is not the case for the pairs of weakly expressed or silent convergent genes and DGPs (regardless of their expression level) that do not tend to form contact domains and stripes (Extended Fig. 6).

**Figure 6.**
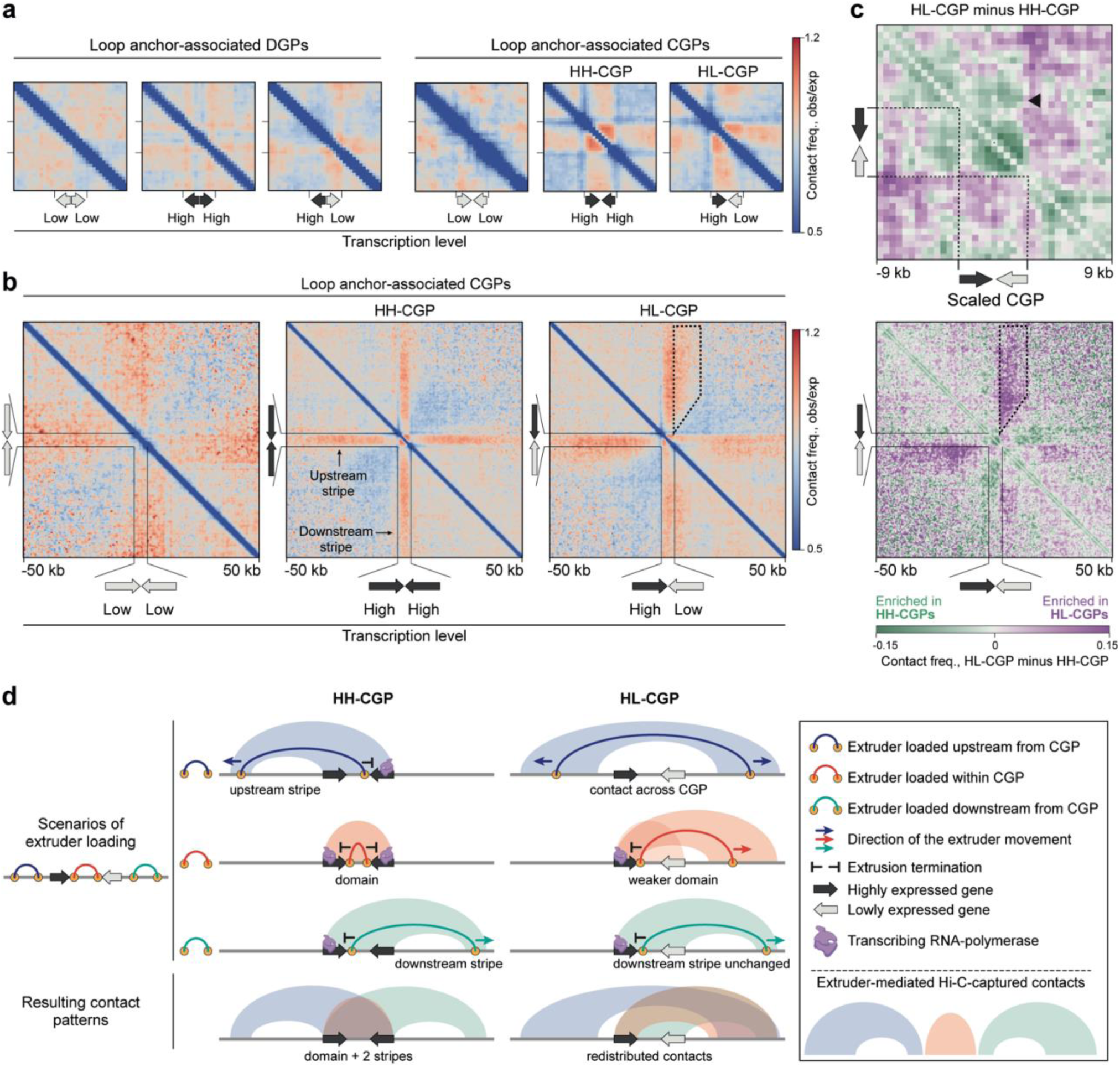
Convergent gene pairs shape chromatin interactions. **a**, Averaged Hi-C maps centered at convergent and divergent gene pairs with different levels of transcription. These pictures show the contact patterns within a gene pair and in its immediate vicinity. **b**, Averaged Hi-C maps centered at convergent gene pairs with different levels of transcription (genes transcribed at high and low levels are highlighted with dark gray and light gray, respectively). The area where contacts are enriched in HL-CGP is demarcated with a dashed line. **c,** Subtraction of averaged Hi-C maps of scaled CGPs. The black triangle in the upper panel shows decreased contacts between the weakly expressed 3’-gene and regions upstream of the CGP. In the bottom panel, the area where contacts are enriched in HL-CGP is demarcated with a dashed line. **d**, Schematic representation of contact patterns generated by interactions between extruders and RNA-polymerases at CGPs.

We then assessed the impact of imbalanced transcription in CGPs and considered pairs where one gene is highly transcribed, and the other is lowly transcribed (‘high-low CGPs’, HL-CGPs). We averaged the Hi-C maps around HL-CGPs with a highly transcribed gene always placed at the left, and a lowly transcribed one at the right. This revealed a partial and, most importantly, one-sided degradation of the CGP contact domain with the loss of short-range contacts, specifically at the side of the lowly expressed gene (Fig. 6c upper panel). This suggests that the extruder bypassesthe lowly gene boundary. In line with that, the lowly expressed gene loses contacts within the ∼50 -kb vicinity upstream of HL-CGP (from the side of the highly transcribed gene; Fig. 6c). We also observed an asymmetrical gain of contacts across HL-CGP, with the immediate ∼15-kb downstream flank of the HL-CGP increasing interactions with ∼50-kb region upstream of the HL-CGP (Fig. 6c lower panel, dashed area).

All these observations could be explained in terms of interactions of the extruders loaded within and outside CGPs with transcribing RNA-polymerase. Extruders, halted within CGPs, compact it into a self-interacting domain (Fig. 6d, HH-CGP, upper middle line). Extruders, entrapped at CGPs from one side, and continuing uni-directional extrusion from another side (and thus moving away from the CGP) produce stripes (Fig. 6d, HH-CGP, uppermost and lowermost lines). Redistribution of contacts across HL-CGP might be caused by the weak barrier activity of the lowly transcribed gene for the extruders coming from the loci upstream of the CGP. These extruders successfully pass the CGP, thus weakening the contact domain at CGPs and mediating gain of contacts between the CGP flanks (Fig. 6f, HL-CGP).

Thus, CGPs shape chromatin looping by an interplay between extrusion and transcription. In the Hi-C map, this is manifested in the formation of contact domains and stripes at short and long genomic ranges, respectively.

### An interplay between extrusion and transcription at Dicty loop anchors

To get further confirmation on our hypothesis that the chromatin looping in Dicty can be explained by an interplay between loop extrusion and convergent gene transcription polymer simulations were performed (Fig. 7a). Chromatin is represented as a standard bead-spring model of a polymer chain, with an extruder loading at an arbitrary monomer. The extruder contains two *molecular motors*, each of them reels in DNA and stalls independently, resulting in two-sided loop extrusion^72^ (see Supplementary Methods for more details and^73^ for a comprehensive review). We assumed that either another cohesin or transcribing polymerase acts as a barrier to the moving extruder. Additionally we assumed that the transcribing polymerase can push the extruder co - directionally with its movement.

**Figure 7.**
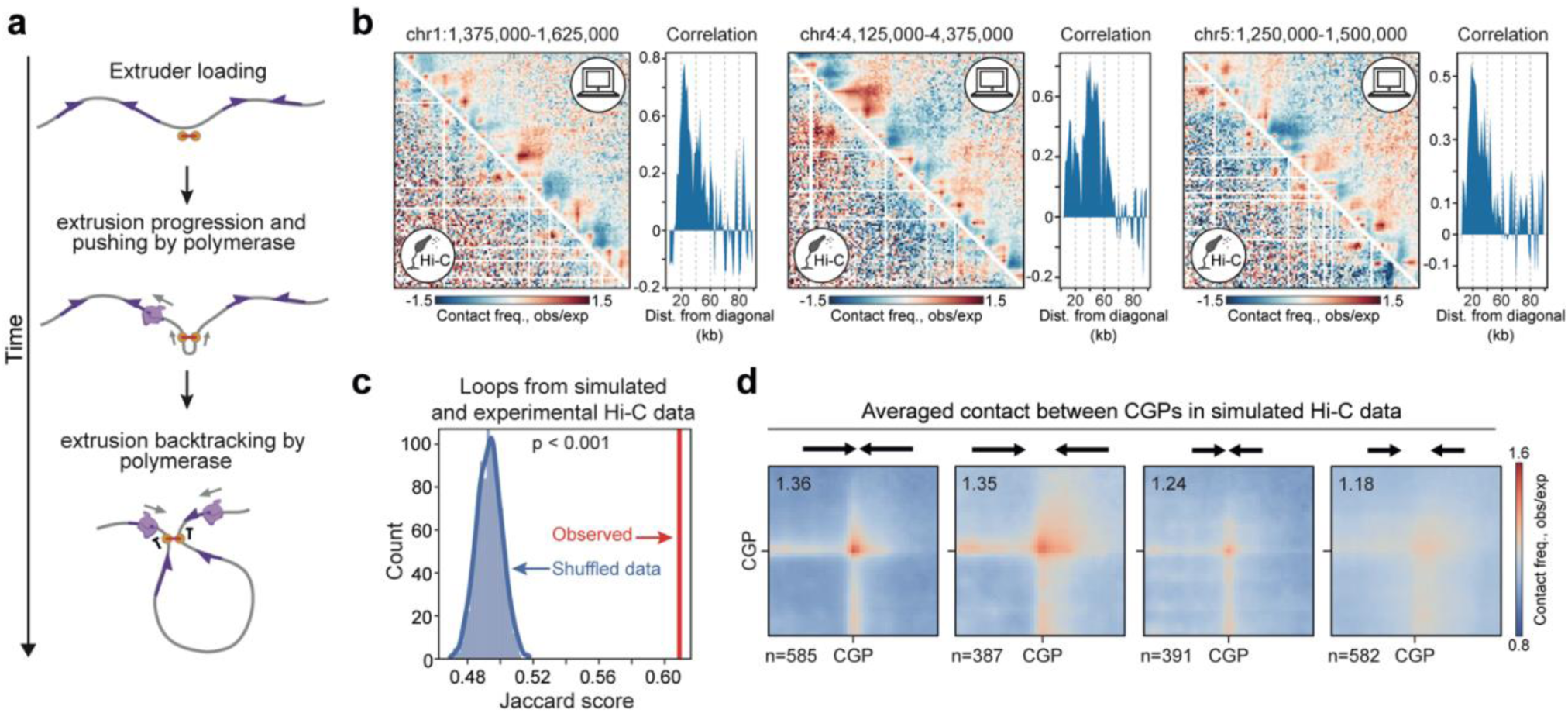
Simulations of Dicty chromatin folding based on interplay between extrusion and transcription. **a**, A scheme of extrusion progression and termination at CGPs as a result of collision with transcribing RNA-polymerase. **b,** Representative examples of simulated Hi-C data (upper map) in comparison with experimental Hi-C data (bottom map). Distributions of the Pearson correlation coefficient averaged over different genomic distances are shown to the right of the maps. **c,** Jaccard score for the loops (taken as intervals) in simulated and experimental Hi-C data. Permutation test p-value is shown. **d,** Averaged contacts between loop anchor-associated CGPs in simulated Hi-C data (compare with Figure 5f which demonstrates averaged contacts between CGPs in experimental Hi-C data). Schematic representation of CGP composition and the numbers of CGPs and are shown above and below the heatmaps, respectively.

To define the parameters of transcription and extrusion the following observations were taken into consideration. Firstly, the RNA polymerase speed has been previously assessed for Dicty to be 1.3 kb/min^74^, similar to that of *Drosophila* (∼1 kb/min)^75^ and mammals (1.25 kb/min-3.5 kb/min)^76^, but faster than that of the yeast RNA polymerase (∼0.8 kb/min)^77^. Secondly, using yeast data on the RNA polymerase density^78^, we have assumed that there is approximately one RNA polymerase molecule per 17.4 kb of the genome with the expected average gene initiation rate of 0.115 (see Supplementary Methods and Supplementary Table 9). Finally, we have varied the extrusion speed, as some studies suggest that it is slower than the speed of transcribing RNA polymerase^79^, whereas other papers claim that extruders move significantly faster^32^.

We used these estimates as a starting point for the parameter sweep. First, we performed preliminary optimization of the transcription initiation rate and the extruder/polymerase speed ratio in a model polymer containing four CGPs. This optimization was done to minimize the difference of loop strength between simulated and observed data. We used the median strength of contacts between neighboring CGPs in the Hi-C data for vegetative cells as a target value, and measured the loop strength between CGPs in the model polymer to determine suitable parameter windows (Extended Fig. 7a). Relatively narrow ranges of both the transcription initiation rate and the extruder/polymerase speed ratio appear to be optimal: the extruder/polymerase speed ratio being equal to 0.05 −0.1 and the transcription initiation rate being equal to 0.05-0.1 (Extended Fig. 7b). The latter is close to *in vivo* measurements in Dicty^78^.

Next, we performed local optimization of parameters using a random genomic locus and the Pearson correlation between simulated and experimental region on Hi-C maps as a measure of simulation performance. With this metric, we found that the following set of parameters appear to be optimal: contact radius is 1.0-1.2 at the transcription initiation rate of 0.05 (Extended Fig. 7c, left panel), while the extruder/polymerase speed ratio is 0.1 (Extended Fig. 7c, right panel). Both sets of parameters indicated that the extruder processivity is 30 kb, which is close to the average loop size in Dicty (Fig. 2b), but quite low compared to the 100-1,000 kb extruder processivity in yeast and mammals^68,72,80^.

Finally, we proceed to whole-genome simulations, and confirmed that the extruder processivity of 30 kb (Extended Fig. 8a) and the extruder/polymerase speed ratio of 0.1 (Extended Fig. 8b) resulted in a higher overall correlation between the experimental and simulated data. At the same time, we observed a substantial variability in the quality of predictions between individual loci (Extended Fig. 8c, d). Notably, the extruder/polymerase speed ratio of 0.1 corresponds to the absolute extrusion speed of 40 bp/sec. This is substantially slower than the proposed speed for other species, but close to the estimate of the extrusion speed of 100 bp/sec obtained from the estimations of extruder lifetime and the time of chromatin attachment^79^.

The resulting simulations demonstrate close similarity to the experimental Hi-C data (Fig. 7b), confirming our hypothesis of loop extrusion playing a major role in the organization of chromatin in Dicty through an interplay with transcribing polymerases. To test whether loop patterns are generally reproduced in simulations genome-wide, we called loops on a simulated Hi-C map using Сhromosight tool^49^ with the same parameters as for the experimental data, which resulted with 1453 loops. The Jaccard index for two loop sets is 0.61, which is significantly higher than for random regions (Fig. 7c, *p* < 0.001, permutation test). To further check the consistency between simulations and experimental data, we plotted the average loop between CGPs from Figure 5f using the simulation data. The strongest contacts between the long-gene and short-spacer CGPs were reproduced in simulations (Fig. 7d). We thus concluded that the Dicty genome loops could be explained by relatively slow extruders with low processivity - most likely cohesins - terminating at convergent pairs of highly transcribed genes.

### Potentially decreased ATPase activity of Dicty cohesin

To understand the possible reasons for low extrusion speed and processivity of Dicty cohesin, we performed a comparative analysis of amino acid composition of Dicty cohesin subunits with orthologs from 18 species selected based on iTOL eumetazoa branches using protein sequences from the UniProt database (Extended Fig. 9a, b). ATP binding is required for cohesin loading onto DNA^81^, and ATP hydrolysis is required for the extrusion progression^73^. We hypothesized that the low extrusion speed and/or processivity could be caused by the less efficient ATP binding and/or hydrolysis. Thus, we focused research on the SMC3 and SMC1 subunits, as they form a heterodimeric ATP-binding cassette (ABC) with two ATPase sites within the cohesin’s head, each comprising the Walker A and Walker B motifs, D-loop, and Signature motif (Fig. 8a).

**Figure 8.**
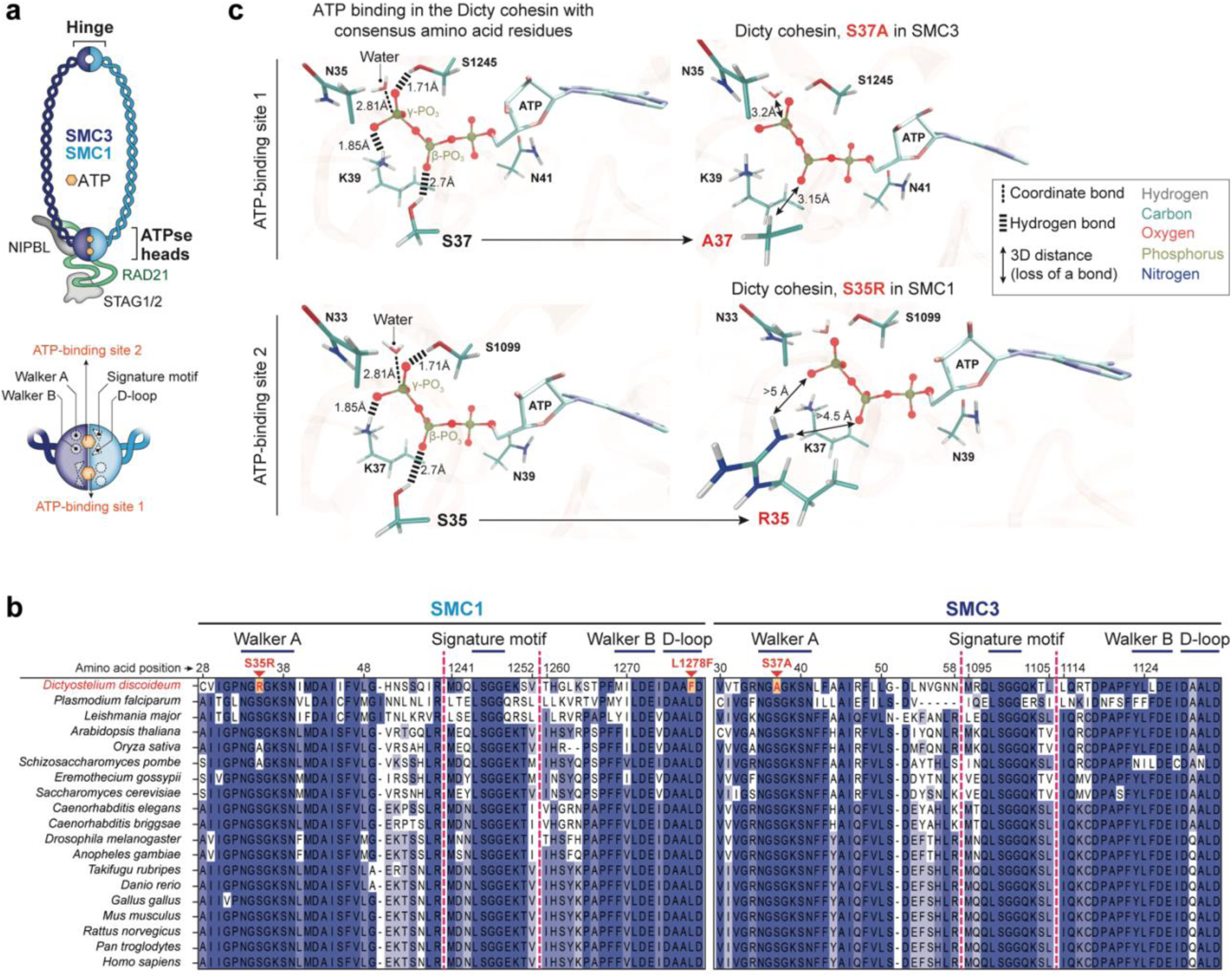
Molecular modeling of ATP binding in ATPase heads of the Dicty cohesin. **a**, General scheme of the cohesin complex structure. Positions of ATP-binding motifs in ATPase heads are shown. **b,** Alignments of amino acid sequences of SMC1 and SMC3 orthologs from different organisms. Substitutions in Dicty cohesin are shown in red. Dashed magenta lines show the positions of skipped parts of the alignment. Amino acid positions are shown according to Dicty SMC1 and SMC3. **c,** Effects of amino acid substitutions in Walker A motifs on ATP binding. Left panels: ATP binding by Walker A motifs in the Dicty cohesin with consensus serine residues in positions 35 (in SMC1) and 37 (SMC3) in the Walker A motif. In sites 1 and 2, ATP is coordinated by the Walker A motif residues from SMC1 and SMC3, respectively. Only bonds affected by the S35R and S37A substitutions are shown. Right panels: consensus S35 and S37 are respectively replaced with arginine (R35) and alanine (A37) residues, originally present in the Dicty cohesin. K37 and K39 are the lysine residues in the Walker A motif of the SMC1 and SMC3 subunits, respectively. γ-PO_3_ and β-PO_3_ - gamma- and beta-phosphate group in ATP.

As expected^82^, protein sequence alignment using MUSCLE revealed a high overall conservation of the ABC-like ATPase domains among the selected organisms (Fig. 8b). However, we observed several amino acid substitutions in Dicty SMC3 and SMC1 orthologs, namely S37A (SMC3); S35R and L1278F (SMC1), that distinguish them from all other organisms from the list. Notably, the K38A substitution in human SMC3 has been shown to significantly impair cohesin ATPase activity and DNA binding^71^. In Dicty SMC3 K38A corresponds to K39 which is in proximity to Dicty-specific A37. Additionally, L1278F in Dicty SMC1 corresponds to the L1166N substitution in *Schizosaccharomyces pombe* SMC1 that leads to altered ATPase activity^83^.

To further test the effects of these substitutions on ATP binding and hydrolysis, we estimated the induced changes in nucleotide binding energy (*ΔΔG*_*bind*_) (Extended Fig. 9c) and the conformational mobility. Our analysis revealed that the S37A substitution in the SMC3 Walker A domain and the S35R substitution in the SMC1 Walker A domain destabilize ATP binding (Fig. 8c). Specifically, S37A increases *ΔG*_*bind*_ by 0.8 ± 0.2 kcal/mol (which is equivalent to 1.3 kT) due to the loss of a long hydrogen bond (2.70 Å) between the serine hydroxyl group and one of the β-phosphate oxygens. However, it does not significantly disturb the conformational dynamics of ATP in its binding site.

S35R substitution in the SMC1 Walker A domain affects the serine residue, homologous to S37 in the SMC3, and is responsible for maintaining β-phosphate as well. Due to its relatively large side chain and positive charge, arginine experiences strong repulsion from nearby residues (especially from K37 and N33, see Fig. 8c) and turned out to be unable to form a stable short hydrogen bond with the γ- or β-phosphate oxygens. This leads to an increase in *ΔG*_*bind*_ by 1.3 ± 0.3 kcal/mol (which is equivalent to 2.2 kT). Two remaining substitutions, L1278F and V1270I, showed no significant impact on ATP binding and hydrolysis efficacy, even in a cooperative manner. Taken together, these observations suggest that Dicty cohesin might have a low extrusion speed and/or low processivity, inline with our conclusions from polymer simulations.

## Discussion

Our findings suggest that Dicty chromatin is predominantly organized into consecutive, non-hierarchical loops, whose interior is only weakly insulated from the adjacent loci (Fig. 2с). Loop anchors are sites of convergent gene orientation (Fig. 4e), which partition chromosomes into transcription-facilitating neighborhoods. Genes, located within the same loop, have similar expression trajectories (Fig. 3b, c) during development and are often functionally-related (Fig. 3d), suggesting that at least some loops may constitute functional domains of the Dicty genome.

Based on the experimental data and the results of polymer simulations, we propose that loop anchors are established by an interplay between transcription and extrusion at CGPs. RNA polymerases are known to act as moving barriers for the cohesin- and condensin-driven extrusion^32,38^. The question is: how oncoming and co-directional transcription (relative to the extrusion direction) contributes to the CGP barrier activity (Fig. 6c)? In this respect, the important observation is that HH-CGPs located in loop anchors form contact stripes (Fig. 6b, d). This suggests that such gene pairs constitute a barrier for extruders coming from the outside from both directions, and, hence, they mediate loop formation. Comparison of HH-CGPs with HL-CGPs revealed the loss of stripe selectively at the side of the weakly transcribed gene (right gene in Fig. 6c, d), suggesting that stripes are formed by head-to-tail collisions of extruder and RNA polymerase. The increased contact frequency between loci located immediately downstream from the weakly transcribed gene and regions upstream of the HL-CGP (Fig. 6c, upper panel, Fig. 6d) implies, that extruders coming from the upstream pass HL-CGPs more efficiently, if compared to HH-CGPs, because they do not face oncoming transcription of the right gene. At the same time, insulation at HL-CGPs is generally preserved, indicating that active transcription of the left gene (which is oncoming for extruders moving from the downstream direction) constitutes an effective barrier for the extrusion. In this regard, CGP may be described as a diode barrier for the extruder complexes. The directional barrier arises due to the difference in transcription levels of the genes that make up the convergent pair, which ensures effective extruder passage primarily from the side of the highly transcribed gene, possibly because co-directional transcription can be a less effective barrier than oncoming one^32^.

Interestingly, the extruders passing through HL-CGP from the upstream loci (that is, from the side of the highly transcribed gene) mediate an increase of contact probability over a relatively small range of genomic distances (5-10 kb, dashed area in Fig. 6c lower panel). This further confirms low processivity of the Dicty cohesin, which is in agreement with the results of polymer simulations. Moreover, the molecular dynamics simulation demonstrates that several amino acid substitutions that distinguish the Dicty cohesin ABC-like ATPse domains from these domains of other cohesins may decrease the rate of ATP hydrolysis. This would affect the extrusion parameters, probably decreasing its speed and/or processivity, and reduce the probability of passing through the barriers such as single RNA polymerase molecules^32^. Low cohesin processivity may also deplete intra-loop interactions, thus preventing folding of a loop into a contact domain (Fig. 2d), and contribute to the absence of loop hierarchy. In mammals, the increase in cohesin processivity upon depletion of the release factor WAPL results in a more hierarchical loop profile^84^. Also, the absence of loop hierarchy in Hi-C maps suggests that only a subset of loops is formed at a time in each individual Dicty cell, further pointing to low efficiency of the extrusion process.

Why do neither active DGPs nor stand-alone highly transcribed genes act as barriers for the extrusion? One possibility is that not only transcription *per se*, but some other factors contribute to the cohesin stalling. A crucial difference between CGPs and DGPs is the type of supercoiling generated by elongating RNA polymerases: CGPs and DGPs accumulate positive and negative supercoils, respectively^85^. A number of observations indicate that supercoiling is tightly linked to the activity of SMC complexes. It has been recently shown that two other SMC-complexes – condensin^86^ and Smc5/6^87^ from budding yeast – bind the tips of supercoiled DNA plectonemes and start extrusion. In yeast, CGPs with short intergenic spacers accumulate high levels of positive supercoiling^88^. The yeast cohesin is also reported to preferentially compact positively supercoiled DNA^89^. Besides facilitating SMC-complex binding, supercoiling may potentially act as a barrier for the extrusion^90^. Since loops in the Dicty chromatin are predominantly formed between CGPs of long, highly expressed genes with short spacers (Fig. 8c), we suppose that positive supercoiling generated within these CGPs may contribute to the termination of extrusion. Further, supercoils *per se* can compact chromatin, and it has been proposed as a mechanism of the contact-domain formation in the dinoflagellate genome^33^. Since we observed that loop anchor-associated CGPs are organized into contact domains whose structure depends on transcription level of both genes in a pair (Fig. 6c, d), it is tempting to assume that Dicty cohesin complexes loaded at supercoiled regions (most probably, at intergenic spacers) within the CGP collide with RNA-polymerases. This results in CGP compaction into the contact domain by halted extruders.

Finally, we should note that while we propose cohesin-driven extrusion as a mechanism of loop formation in the Dicty chromatin, we cannot rule out the contribution of other SMC complexes.

## Online Methods

### Dictyostelium discoideum cultivation and storage

*Dictyostelium discoideum* AX4 strain was obtained from the dictyBase Stock Center (strain ID: DBS0237637). Culture was started from the frozen stock on SM agar plates (1 g of anhydrous glucose, 1 g of Proteose Peptone A (Lab M; #MC011), 0.1 g of yeast extract, 1.5 g of agar, 8.62 10× KK2 buffer, MilliQ water to 100 ml; after autoclaving, 415 μl of 1M MgSO_4_ were added^91^). 10× KK2 buffer composition (for 50 ml): 1.1 g of KH_2_PO_4_, 0.458 g of K_2_HPO_4_×3H_2_O, MilliQ water to 50 ml. Frozen stock was thawed and mixed with 10 ml of the modified HL5 axenic growth medium ^92^ in a 100-mm plastic Petri dish (2.5 g of Tryptose (Lab M; #MC008), 2.5 g of Proteose Peptone A (Lab M; #MC011), 2.5 g of yeast extract, 5 g of anhydrous glucose, 1.27 ml of 1M KH_2_PO_4_, 0.65 ml of 1M Na_2_HPO_4_, MilliQ water to 500 ml; note that in our experience, not all Tryptoses and Peptones are suitable for the *Dictyostelium* axinic growth). The dish was incubated for 30 minutes at 20°C. The medium was removed, and attached cells were rinsed off from the dish by pipetting in 10 ml of fresh HL5 medium. Cells were harvested for 5 minutes at 400 g (room temperature; hereinafter 20°C). 100 μl of the mixture of fresh overnight culture of *E. coli* DH5α and 0.5-2×10^6^ *Dictyostelium* cells was spread on 100-mm SM agar plates using a sterile glass spatula. Plates were inverted, placed in a humid chamber (plastic bag with wet filter paper) and incubated at 20°C for 5-7 days. After the appearance of growth plaques, cells were scraped using a bacteriological loop and inoculated in 100 ml of modified HL5 axenic growth medium supplemented with 1× penicillin/streptomycin (Thermo Fisher Scientific; #15140122) or kanamycin (Thermo Fisher Scientific; #15160054) in a 500-ml glass flask and incubated at 20°C with shaking at 200 rpm. Cultures were maintained at a density of 0.2-1×10^6^ cells per ml for no more than 3 weeks. For long-term storage, cells from fresh culture were harvested for 5 minutes at 400 g (4°C) and resuspended in ice-cold HL5 medium supplemented with 5% of dimethylsulfoxide (DMSO, Sigma-Aldrich; #D8418) to a concentration of 5×10^6^ cells per ml. The suspension was aliquoted into 1.8-ml cryovials, incubated at −80°C for 12 hours in an alcohol-free cell freezing container and stored in liquid nitrogen.

### Multicellular development

Induction of multicellular development was performed by starvation on KK2 agar plates according to previously published protocol^91^ with several modifications. 3×10^8^ of cells from fresh axinic culture in HL5 medium (1-2 days, 0.5-1×10^6^ cells per ml) were harvested for 10 minutes at 400 g (room temperature), washed once with 50 ml of ice-cold 1× KK2 buffer and once with 50 ml of ice-cold Development buffer (5 mM Na_2_HPO_4_, 5 mM KH_2_PO_4_, 1 mM CaCl_2_, 2 mM MgCl_2_). The pellet was resuspended in 1 ml of Development buffer. The suspension was spreaded on 150-mm Petri dishes with 1.5% agar prepared on 1× KK2 buffer. Dishes were wrapped with the Parafilm, placed in a humid chamber and incubated at 22°C for 2 (starvation), 5 (early aggregation) and 8 hours (late aggregation). The development progress accompanied by the appearance of characteristic spiral-like patterns formed by migrating cells has been monitored by visual inspection of the cell layer at the agar surface. At each time point, cells were rinsed off from the agar surface by pipetting in 10 ml of ice-cold 1× KK2 buffer. Cells were then harvested for 5 minutes at 400 g (4°C). Pellet was resuspended in 10 ml of ice-cold 1× KK2 buffer, the suspension was splitted into two aliquots in a ratio of 1:4, and cells were harvested as described above. The cell pellet from the smaller aliquot was snap-frozen in a liquid nitrogen for the subsequent RNA isolation. The cell pellet from the larger part was resuspended in 50 ml of 1× KK2 buffer and used for the Hi-C library preparation.

### Hi-C library preparation

Hi-C libraries were prepared as described in^93^ with several modifications. Cells were cross-linked in 50 ml of 1× KK2 buffer supplemented with fresh 2% formaldehyde (Sigma-Aldrich; #F8775) for 10 minutes at room temperature. Excess of formaldehyde was quenched with 125mM glycine for 5 min. Cells were centrifuged (1,000 g, 10 minutes, 4°C), resuspended in 100 μl of 1× PBS, snap-frozen in liquid nitrogen, and stored at −150°C. Defrozen cells were lysed in 1.5 ml of Isotonic buffer (50 mM Tris-HCl pH 8.0, 150 mM NaCl, 0.5 % (v/v) NP-40 substitute, 1 % (v/v) Triton X-100, 1× Protease Inhibitor Cocktail (Bimake; #B14001)) on liquid ice for 15 minutes. Cells were centrifuged at 5,000 g for 7 minutes, resuspended in 100 μl of 1.12× DpnII buffer (NEB; #B0543S), and pelleted again. The pellet was resuspended in 200 μl of 0.1% SDS in 1.12× DpnII buffer and incubated at 65°C for 10 minutes without shaking. Then, 330 μl of 1.12× DpnII buffer and 52.7 μl of 20% Triton X-100 were added, and the suspension was incubated at 37°C for 1 hour with shaking (1,400 rpm).

Next, 600 U of DpnII restriction enzyme (NEB; #R0543M) were added, and the chromatin was digested overnight (14-16 hours) at 37°C with shaking (1,400 rpm). On the following day, 200 U of DpnII restriction enzyme were added, and the cells were incubated for 2 hours. DpnII was then inactivated by incubation at 65°C for 20 minutes. After DpnII inactivation, the nuclei were harvested for 10 minutes at 5,000 g in a pre-cooled centrifuge (4°C), washed twice with 200 μl of 1,2× NEBuffer 2.1 (NEB; #B7202S), and resuspended in 125 μl of 1,2× NEBuffer 2.1. Cohesive DNA ends were biotinylated by the addition of 25 μl of the biotin fill-in mixture (0.025 mM dCTP (Thermo Fisher Scientific; #R0151), 0.025 mM dGTP (Thermo Fisher Scientific; #R0161), 0.025 mM dTTP (Thermo Fisher Scientific; #R0171), 0.025 mM biotin-14-dATP (Thermo Fisher Scientific; #19524-016), and 0.8 U/μl Klenow enzyme (NEB; #M0210L)). The samples were incubated at 37°C for 90 minutes with shaking (900 rpm). Nuclei were pelleted at 5,000 for 5 minutes, resuspended in 100 μl of 1× T4 DNA ligase buffer (Thermo Fisher Scientific; #EL0011), and pelleted again. The pellet was resuspended in 400 μl of 1× T4 DNA ligase buffer, and 50 U of T4 DNA ligase (Thermo Fisher Scientific; #EL0011) were added. Chromatin fragments were ligated at 22°C for 6 hours with shaking (1,400 rpm). The cross-links were reversed by overnight incubation at 65°C in the presence of Proteinase K (100 μg/ml; Sigma-Aldrich; #P2308) and 1% of SDS. After cross-link reversal, the DNA was purified by single phenol-chloroform extraction followed by ethanol precipitation with 20 μg/ml of glycogen (Thermo Scientific Scientific; #R0561) as a co-precipitator. After precipitation, the pellets were dissolved in 100 μl of 10 mM Tris-HCl (pH8.0). To remove residual RNA, samples were treated with 20 μg of RNase A (Thermo Fisher Scientific; #R1253) for 45 minutes at 37°C. To remove residual salts and DTT, the DNA was additionally purified using Agencourt AMPure XP beads (Beckman Coulter; #A63881). Biotinylated nucleotides from the non-ligated DNA ends were removed by incubating the Hi-C libraries (2 μg) in the presence of 6 U of T4 DNA polymerase (NEB; #M0203L) in NEBuffer 2.1 supplied with 0.025 mM dATP (Thermo Fisher Scientific; #R0141) and 0.025 mM dGTP at 20 °C for 4 hours. Next, the DNA was purified using Agencourt AMPure XP beads. The DNA was then dissolved in 500 μl of sonication buffer (50 mM Tris-HCl (pH8.0), 10 mM EDTA, 0.1 % SDS) and sheared to a size of approximately 100 - 500 bp using a VirSonic 100 (VerTis). The samples were concentrated (and simultaneously purified) using AMICON Ultra Centrifugal Filter Units (Millipore; #UFC503096) to a total volume of approximately 50 μl. The DNA ends were repaired by adding 62.5 μl of MilliQ water, 14 μl of 10× T4 DNA ligase reaction buffer, 3.5 μl of 10mM dNTP mix (Thermo Fisher Scientific; #R0191), 5 μl of 3 U/μl T4 DNA polymerase, 5 μl of 10 U/μl T4 polynucleotide kinase (NEB; #M0201L), 1 μl of 5 U/μl Klenow DNA polymerase, and then incubating at 25°C for 30 min. The DNA was purified with Agencourt AMPure XP beads and eluted with 60 μl of 10mM Tris-HCl (pH 8.0). To perform an A-tailing reaction, the DNA samples were supplemented with 7.5 μl of 10× NEBuffer 2.1, 1.5 μl of 10mM dATP, 1,5 μl of MilliQ water, and 4.5 μl of 5 U/μl Klenow (exo-) (NEB; #M0212S). The reactions were carried out for 30 minutes at 37°C in a PCR machine, and the enzyme was then heat-inactivated by incubation at 65°C for 20 minutes. The DNA was purified using Agencourt AMPure XP beads and eluted with 100 μl of 10mM Tris-HCl (pH8.0). Biotin pulldown of the ligation junctions: 10 μl of MyOne Dynabeads Streptavidin C1 (Thermo Fisher Scientific; #65001) beads washed 2 times with the TWB buffer (5 mM Tris-HCl (pH8.0), 0.5 mM EDTA (pH8.0), 1 M NaCl, 0.05 % Tween-20), resuspended in 200 μl of 2× Binding buffer (10 mM Tris-HCl (pH8.0), 1 mM EDTA, 2 M NaCl) and added to 200 μl of DNA. Biotin pulldown was performed for 30 minutes at 25 °C with shaking. Next, beads with captured ligation junctions were washed once with 1× Binding buffer, once with 1× T4 DNA ligase buffer, and then resuspended in 50 μl of adapter ligation mixture comprising of 41.5 μl of MilliQ water, 5 μl of 10× T4 DNA ligase reaction buffer, 2.5 μl of Illumina TruSeq adapters, and 1 μl of 5 U/μl T4 DNA ligase. Adapter ligation was performed at 22 °C for 2.5 hours, and the beads were sequentially washed twice with 200 μl of TWB, twice with 200 μl of CWB (10 mM Tris-HCl (pH8.0) and 50 mM NaCl), and then resuspended in 25 μl of MilliQ water. Test PCR reactions containing 4 μl of the streptavidin-bound Hi-C library were performed to determine the optimal number of PCR cycles required to generate an amount of PCR products sufficient for sequencing. The PCR reactions were performed using KAPA High Fidelity DNA Polymerase (Kapa Biosystems; #08201595001) and Illumina PE1.0 and PE2.0 PCR primers (10 pmol each). The temperature profile was 5 minutes at 95°C, followed by 6, 9, 12, 15, and 18 cycles of 20 sec at 98°C, 15 sec at 65°C, and 20 ses at 72°C. The PCR reactions were separated on a 2 % agarose gel containing ethidium bromide, and the number of PCR cycles necessary to obtain a sufficient amount of DNA was determined based on the visual inspection of gels (typically 8-12 cycles). Four preparative PCR reactions were performed for each sample. The PCR mixtures were combined, and the products were purified with Agencourt AMPure XP beads. Libraries were sequenced on an Illumina Novaseq 6000 by 100-bp paired-end reads.

### RNA-seq library preparation

10^8^ of cells from fresh culture (1-2 days, 0.5-1×10^6^ cells per ml) or from development plates were harvested for 5 minutes at 400 g (room temperature), washed with 5 ml of 1× KK2 buffer and snap-frozen in a liquid nitrogen. RNA extraction was carried out using an RNeasy Mini kit (Qiagen) following the manufacturer’s instructions. RNA quality was assessed using capillary electrophoresis with a Bioanalyzer 2100 (Agilent). For library preparation, a TruSeq RNA Sample Prep kit v2 (Illumina) was used following the manufacturer’s instructions. After preparation, libraries were quantified using a Qubit fluorometer and quantitative PCR and sequenced with HiSeq 2000 with read lengths of 51 nt.

### Hi-C analysis

#### Raw read processing

Hi-C paired-end reads were processed with distiller-nf pipeline version 0.3.3 with default parameters. In summary, the reads were initially mapped to *Dictyostelium discoideum* reference genome using bwa mem (0.7.17-r1188). Subsequently, the mapped reads were processed into pairs of aligned reads using pairtools (0.3.1-dev.1) with walks-policy “mask”. Only reads mapped with a mapping quality (MAPQ) of 30 or higher were considered as potential contact pairs. Multimappers, one-sided mapped reads, and potential PCR duplicates were filtered out. The resulting set of pairs was converted into 100-binned contact matrices using cooler (0.8.7).

#### Correlation Calculation

To evaluate the reproducibility of Hi-C replicates, correlation coefficients were calculated using the Python implementation of HiCRep v0.2.6. This process involved generating a correlation matrix, where each cell represents the correlation between two Hi-C samples. Subsequently, the correlation matrix was visualized as a heatmap, with annotations indicating the HiCRep coefficient values. The heatmap visualization is available in Figure S1b.

#### Subsampling and merging replicates

The impact of read depth on feature calling is well-documented^94^. Given that and the high correlation between replicates, we opted to merge them. Specifically, we retained reads from the six main scaffolds, subsampled them to approximately 22 million reads each, and merged them with samples from the corresponding developmental stage.

#### Scaling plot

The scaling plot for Figure 1h was constructed using the expected_cis function from cooltools v0.5.4. Parameters used for this analysis include ignore_diags=2, smooth=True, aggregate_smoothed=True. The plot was generated in log-log coordinates to better visualize the relationship between genomic distance and average contact frequency.

#### Compartments

The genome-wide eigenvector for Figure S1d was computed at a resolution of 100 kb resolution and phased with GC-content using the eigs_cis function from cooltools v0.5.4.

#### Average Rabl configuration

The average Rabl heatmap represents the mean frequencies of interchromosomal interactions, emphasizing chromatin contacts enriched due to the Rabl configuration of chromosomes. To generate Figure 1c, sampled and filtered Hi-C maps at 100 kb resolution were subjected to an iterative correction procedure using cooler software (v0.8.6)^95^. The following command was applied: “cooler balance --trans-only --min-nnz 300”. The matrices of interchromosomal contacts were extracted from the resulting maps and rotated so that the centromere appeared near the upper left corner. Chromosome 2 was excluded from the further analysis due to a unique large loop that interferes with the Rabl structure. Next, each rotated matrix was normalized by its mean value with NaN values ignored and rescaled to a common size of 25×25 using the “lib.numutils.zoom_array()” function from the cooltools package (v0.4.1) with the parameters “same_sum=False, order=1”. The resulting matrices of trans interactions were superimposed one upon the other, and the mean value for each pixel was calculated again ignoring NaN values, Finally, the resulting matrix was log2-transformed.

#### Las-loops

Due to the relatively low number, las-loops were annotated manually at 4 kb resolution using Higlass.

#### Long non-coding RNAs

The list of long non-coding RNAs (lncRNAs) in Dicty was obtained from^21^. The count matrix for genes was sourced from^58^, and it was TPM-normalized. For Figure 1f delta was computed as the differences between the expression levels at 8 hours and 0 hours after starvation induction. In Figure 1e, lncRNAs were assigned to be anchor-associated if their gene intersected with any las-loop anchor.

#### Loop annotation using LASCA

Dicty Hi-C maps in cool format with 2000 bp resolution were subjected to analysis using the LASCA pipeline^48^ (https://github.com/ArtemLuzhin/LASCA_pipeline). The following functions and corresponding parameters were applied:

1. *Get_pvalue_v7 (resolution=2000, bin_coverage=0.0, distance_bins=20)*
2. *convert_to_log10_square_matrix*
3. *Get_qvals_mtx_v2(distance_bins=20)*
4. *cluster_dots (min_cluster_siz=3, q_value_treshold=0.01, filter_by_coverage=False)*
5. *Get_coordinates(Intensity=True, as_intervals=True)*

In summary, p-value matrices were generated with *Get_pvalue_v7* function, which were then converted to log10 with the convert_to_log10_square_matrix function. Subsequently, final q-value matrices were generated using *Get_qvals_mtx_v2* function. The coordinates of each loop were determined using the *Get_coordinates* function, with the center being defined as the brighter pixel in the cluster.

#### Loop annotation using Chromosight

Another software of choice utilized in the analysis is chromosight v1.6.3^49^. The following parameters were employed: --min-dist 6000 --max-dist 45000 --pearson 0.2 using subsampled merged Hi-C maps at 2000 bp resolution. Subsequently, the resulting loop set was selected as loops called by both tools, ensuring consistency and reliability in the identified loops.

### Proportion of shared loop anchors

Visual observations confirmed that while loop calling algorithms generally perform well, there is a tendency to make errors when defining loop anchors +/- 1 bin to any side. In order to mitigate these biases and ensure the accuracy of our results, we developed a simple algorithm. This algorithm calculates the fraction of 1-bin loop anchors from any Dicty developmental stage that intersects 3-bin (1 bin offset) or 5-bin (2 bins offset) loop anchors from another stage. The intersections were computed using the *overlap* function from the bioframe v0.4.1.

### Consecutive loops

To distinguish between consecutive and stand-alone loops, we utilized the *closest* function from the bioframe v0.4.1 package. Loops that were located within a distance not exceeding 2 kb (1 bin offset) from any other loop were categorized as consecutive loops. Conversely, loops with a distance greater than 2 kb to the nearest loop were classified as stand-alone loops

### Loop hierarchy plot

To assess loop hierarchy, we only considered loops with a chromosight score greater than

0.2 and a q-value less than 0.05. We then constructed an artificial dataset by pairing the starts of loops with index N and the ends of loops with index N+m, where m ranged from 0 to 3. These pairs were positioned accordingly in Figure 2c.

### Average chromatin feature of different organisms

To compare the average features of different organisms, S-stage *S. cerevisiae* MicroC-XL data from^51^ at a resolution of 1600 bp, spermatogonia-stage Hi-C of *D. melanogaster* from^52^ at a resolution of 5 kb, and in-situ Hi-C of human GM12878 lymphoblastoid from^50^ at a resolution of 5 kb.

### Averaged Hi-C plot

To create an average Hi-C plot, we utilized coolpuppy v1.0.0^96^. All maps were normalized by expected values, which were calculated using the *expected_cis* function from cooltools v0.5.4 with ignore_diags=2. The plotpup module from coolpuppy was employed to generate figures.

For average CGP with different expression levels, additional settings were applied: rescale=True, rescale_size=int(1+flank * 2// resolution), rescale_flank=10, flank=20_000, local=True, flip_negative_strand=True. The flip_negative_strand flag was used to orientate HL-CGPs such that highly transcribed genes are positioned to the left, and weakly transcribed ones to the right of each other.

For average interactions between CGP at given distances, the following settings were added: mindist=500, maxdist = 30,000, by_distance=[5,000, 10,000, 20,000, 30,000], flank=20,000.

### Loop strength

To calculate loop strength, an average was computed in the 3×3 vicinity around the middle pixel in the matrix, normalized by expected values.

### El-loops

To annotate elongated loops and distinguish them from regular loops, the following steps were implemented:

1. Compute the 5’ and 3’ medians by calculating the median of the stack data along the specified dimensions. The 5’ median is computed along the rows within a 5-bin segment with a 2-bin offset to the central pixel, while the 3’ median is computed along the columns within the same 5-bin segment.
2. Calculate the fold-change as the logarithm of the ratio of 5’ track intensity to 3’ track intensity.
3. Classify all loops falling between the 20th and 80th percentiles of fold-change as “regular”, while loops with fold-change values outside this range are categorized as “elongated” loops

### Insulation Score

The insulation score for each genomic bin was calculated by averaging the interaction frequencies between the bin and its neighboring bins within the specified window size. This process enabled quantification of the degree of insulation or boundary strength for each genomic region. Lower insulation scores indicate stronger insulation, suggestive of the presence of boundaries between genomic domains

The insulation score computation using *cooltools insulation* involved several key parameters, including resolution and window size. The genome was partitioned into bins of 2 kb in size. Multiple window sizes were tested, including 6 kb, 10 kb, and 20 kb. After evaluation, the window size of 20 kb was selected for downstream analysis, as it best aligned with visual observations in Higlass.

### Convergence Score

The calculation of the gene convergence profile is as in^60^.

### Convergent gene pair annotation

Convergent gene pairs (CGP) were annotated as two genes in convergent orientations, with each gene being the closest neighbor to the other and situated within a maximum distance of 8 kb from each other.

Divergent gene pairs were annotated as two genes located in divergent orientations, with each gene being the closest neighbor to the other and situated within a maximum distance of 8 kb from each other.

#### Loop anchor-associated convergent gene pair (LA-CGP) annotation

CGP was assigned to a loop anchor if their intergenic is fully located within any loop anchor with 1-bin offset.

#### Gene length groups of CGPs

To separate LA-CGPs into distinct groups, the sum of the lengths of the two genes and the intergenic distance were utilized. A threshold was established based on the medians of these value distributions.

#### Expression groups of CGPs

CGPs were categorized into several groups based on their expression levels, with a low-expression group defined as having less than 2 TPM (Transcripts Per Million) and a high-expression group set with a threshold greater than 8 TPM.

If one gene in the pair was assigned to the low-expression group and the other gene to the high-expression group, then the pair was oriented with the highly expressed gene located to the left and the low-expressed gene located to the right.

### RNA-seq analysis

#### RNA-seq preprocessing

The reads were mapped to the D. discoideum genome (version 2.7) using *hisat2* with default parameters, except for setting the maximum intro length set to 3100 bp. Only unique mapped reads were retained based on the mapq parameter, and duplicates were removed using the *samtools markdup* utility.

For downstream analysis, only genes with at least 1 count in 2 samples were selected. DESeq2 were used for normalization of read counts and estimation of variance.

#### Trajectory gene clusters

First, the count matrix was rlog-transformed using Deseq2. This transformation helps to stabilize the variance across the mean, making the data more suitable for downstream analysis.

Next, the (1 - correlation) matrix was computed for all genes. This matrix reflects the dissimilarity between genes based on their expression profiles across samples.

Subsequently, the k-means algorithm was applied to cluster the genes into 4 clusters, which represent *trajectory gene clusters (TGCs)*. K-means clustering partitions the genes into groups based on similarity in their expression profiles.

Finally, a UMAP plot was generated using the *umap* package for Python v0.5.3.

### Loops groups identified by trajectory gene clusters coverage

Loop clusterization was performed using the k-means algorithm, with the coverage by Trajectory Gene Clusters (TGC) serving as an input data.

### Best Match Average (BMA) Score

The Best Match Average (BMA) score facilitates the assessment of functional similarity between genes, providing a quantitative measure of their biological relatedness^97^. The BMA score between two genes based on their Gene Ontology (GO) term associations was computed using a custom Python script. The score was calculated through the following procedure:

1. Obtain the list of GO terms associated with the two genes.
2. Calculate the Lin similarity score for each pair of GO terms.
3. Compute the maximum similarity score for each GO term in the associations of the two genes.
4. Calculate the Best Match Average (BMA) score by summing the maximum similarity scores for both genes and dividing by the total number of unique GO terms associated with the genes.

### Enrichment analysis

To assess whether a genomic feature (*A*) is non-randomly colocalizes or avoids another feature (*B*), a shuffle (permutation) test was conducted. Here’s a summary of the procedure:

1. The intersection between features *A* and *B* (*A* ∩ *B*) was calculated using bedtools intersect with a fraction overlap threshold of 0.1 (option f=0.1).
2. To perform shuffling, bedtools shuffle was utilized with the option chrom=True to maintain the chromosome structure while randomizing the locations of the features.
3. For each shuffle, the intersection between the shuffled feature *A* and feature *B* (*A*⬚_*s*_ ∩ *B*) was calculated.
4. The number of shuffle iterations was set to 1000 (*N*⬚_*s*_ = 1000).
5. The p-value was computed as the minimum of the ratios 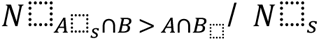 and 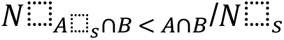, where 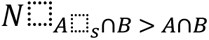 represents the number of shuffles where the intersection between shuffled *A* and *B* is greater than the number of observed intersection *A* ∩ *B*, and 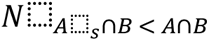 represents the number of shuffles where the intersection is smaller than the observed intersection.

### Blocks of synteny

Synteny blocks between *Dictyostelium discoideum* and *Dictyostelium purpureum* were obtained from^59^.

### Enhancer-like elements annotation

The data utilized in this study was sourced from^20^. Histone modifications, specifically H3K4me1 and H3K27ac ChIP-seq data, underwent preprocessing using the nf-core/chipseq pipeline v1.2.2. Similarly, ATAC-seq was preprocessed using the nf-core/atacseq pipeline v2.0. In cases where peaks from H3K3me1 and H3K27ac intersected, their union was taken. Then this union was intersected with ATAC-seq peaks.

### Coverage Track Generation

The *bamCoverage* tool is employed to generate coverage tracks in BedGraph format from the indexed BAM file. This process involves specifying parameters such as the bin size for resolution, normalization using base pair per million (BPM), applying a smoothing function, centering reads to ensure accurate representation, and providing the effective genome size for proper scaling.

### Housekeeping genes

House-keeping genes were identified using data from^58^ following a modified procedure outlined in^98^ with Dicty-specific modification: (i) genes exhibited expression for all time points; (ii) genes with low variance in expression over time were selected, with a standard-deviation of log2(RPKM) less than 1; (iii) genes with no exceptional expression in any single time point were included. Specifically, genes were excluded if any log-expression value deviated from the averaged log2(RPKM) by two-fold or more. The resulting list contained 1138 genes.

### Logistic regression

Loop anchors were predicted using a logistic regression classifier implemented in scikit-learn library (v.1.0.2). Dicty genome was divided into 2 kb bins which were labeled as either loop anchors or non-anchors. Five bins from the ends of each chromosome were excluded from the analysis. Loop anchor bins were further classified into three categories: insulating/non-insulating anchors of el-loops and anchors of ordinary loops. Since it is possible for a bin to be an anchor of two loops of different types, each bin could belong to more than one category. When fitting the model and predicting loop anchors of one category, bins with loop anchors of other categories were ignored.

Total expression was assigned to a bin as a single feature, representing total RNA-seq expression in the bin adjacent to the left bin and the bin adjacent to the right bin. Convergent expression was assigned to a bin as two features: RNA-seq expression on the plus strand in the bin adjacent to the left bin and expression on the minus strand in the bin adjacent to the right bin. Chromosomes 1, 2, 4, 5 and 6 were used to fit the logistic regression model, and chromosome 3 was used to predict loop anchors.

### Polymer Simulations of 3D Genome

For simulations of loop extrusion, we utilized the classical bead-spring polymer model, implemented through OpenMM^99^ and the polychrom toolkit (https://github.com/open2c/polychrom), to simulate 3D genome contact maps at 250 bp resolution. Similar to previous loop extrusion simulations^32,38,69^, we first sampled trajectories of extruder and polymerase movement on a linear DNA lattice with custom Python code based on previous works. Extruders loaded onto the lattice, moving bidirectionally unless obstructed, while polymerases progressed from gene start to end sites without stalling. Extruders followed specific rules for head-to-head, head-to-tail, and tail-to-head collisions with polymerases. The 1D lattice dynamics served as input for a 3D polymer model, where each extruder created an additional bond and was repositioned to the new location at each 1D step.

First we run simulations of four CGPs with the gene properties (position and orientation) corresponding to the average of the genome. We run the parameter sweep of speed ratio between extruder and polymerase, polymerase transcription rate, *in silico* Hi-C contact distance, and extruder lifetime, and established an initial optimal range of parameters optimizing Hi-C loop strength between neighboring (N) and N+1 CGPs.

We then extended simulations to two 250-Kb regions of the Dicty genome, simulating polymerase and cohesin movement proportional to transcriptional activity. Using RNA-Seq data, we mapped genes to a 1D lattice, assigning relative polymerase initiation rates. Polymer simulations produced *in silico* Hi-C maps, varying key parameters, and comparing to experimental Hi-C maps using Pearson correlation coefficients. Results indicated optimal parameters varied by genomic region, emphasizing the complexity of parameterizing genomic simulations.

For comprehensive whole-genome simulations, we divided the genome into 250-Kb windows, excluding low-mappability regions, and ran polymer simulations for each, varying extruder lifetimes and speed ratios. Globally optimal parameters were identified, producing visually similar *in silico* maps to experimental Hi-C maps, despite regional variations.

For more information on the simulation setup and the parameter sweep, see Supplementary Information.

### Multiple alignment

Protein sequences were obtained from the UniProt database^100^.

Multiple alignment was performed using MUSCLE v3.8.31^101^, and the alignment was visualized using Jalview v2.11.3.2^102^.

### Molecular modeling

The three-dimensional molecular model of Dicty cohesin was constructed using AlphaFold (v.2.3.1, (https://github.com/google-deepmind/alphafold)^103^ in multimer mode with all other parameters set to default values. The final structure consisted of the following segments of domain sequences: 1-178, 1171-1337 of SMC1; 1-165, 1039-1195 of SMC3; 726-809 of Rad21. The most confident model with ipTM+pTM score equal to 0.843 was selected for further analysis.

The positioning of ATP molecules in their binding sites, as well as closest amino acid residues, magnesium ions, and catalytic water molecules, was carefully refined based on available crystalline structures of ATP-cohesin complexes and previous computational investigation of ATP hydrolysis by human cohesin^104–106^. We utilized the CHARMM-GUI web server^107–109^ to manipulate model structure (including the introduction of target amino acid substitutions) and generate molecular mechanics (MM) topologies and input files for molecular dynamics (MD) simulations. The system was dissolved in a cubic 109×109×109 Å^3^ water box and sodium and chloride ions were added to neutralize the solution and adjust NaCl concentration to physiological levels (150 mM). AMBER ff14SB^110^, TIP3P^111^, and GAFF2^112^ force field models were used to parametrize protein, water and ATP molecules respectively.

To investigate the effect of certain amino acid substitutions on the efficacy of ATP binding and hydrolysis, we estimated the change of ligand binding free energy (*ΔΔG*_*bind*_) upon these mutations. For that purpose, we utilized the Quantum Mechanics / Molecular Mechanics – Poisson-Boltzmann Surface Area (QM/MM-PBSA) method^113^. This method, an extension of the classical MM-PBSA^114^, allows us to compute the gas-phase part of the binding free energy at a higher level of theory.

The motivation to employ a hybrid Quantum Mechanics / Molecular Mechanics (QM/MM) approach in this study comes from two primary considerations. Firstly, the complex coordination environment of ATP that, in particular, includes catalytic water molecules and magnesium ion accompanied by its own orientational water molecules poses a challenge for accurate description using purely Molecular Mechanics (MM) force fields. Similarly, the chemical processes occurring within the active site of the protein require a more sophisticated modeling approach. Secondly, the results obtained from quantum mechanical calculations generally offer higher accuracy compared to classical methods. However, it’s worth noting that the utilization of the hybrid methodology forced us to omit the entropic contribution to binding energy due to its poor convergence within short QM/MM MD simulations (0.001-0.01 ns). Nonetheless, we believe its impact on our findings to be minimal within the scope of this study.

We performed all molecular dynamics simulations employing NAMD (v.2.14) software^115^. The typical setup included the usage of Langevin thermostat (T = 300 K), Nosé-Hoover Langevin piston barostat (P = 1 atm), integration step 1 fs and the absence of any bond constraints. The non-bonded cutoff was set to 9 Å, and the long-range electrostatics was computed using particle mesh Ewald (PME) methodology. In the case of QM/MM MD, we utilized the NAMD/ORCA (v.5.0.1) program interface^116^. The QM part was described at the B3LYP/D3BJ/6-31G** level of theory^117^ and consisted of ATP, magnesium ion, catalytic and orientational water molecules, as well as the amino acid residue closest to nucleoside. QM-MM interactions were treated in terms of electrostatic embedding, with Mulliken atomic charges computed for the quantum subsystem at every step of molecular dynamics. We used the link atom scheme to resolve the boundary effects.

Four substitutions were studied: one (*S*35*R*^*SMC*1^) is located within one of two ATP binding cassettes, and two (*S*37*A*^*SMC*3^ and *L*1278*F*^*SMC*1^) within another. *V*1270*I*^*SMC*1^ was relatively far away from both active sites and thus was not included in the QM part of our setup. *ΔΔG*_*bind*_ was calculated as the difference between QM/MM-PBSA binding energies of the variants with conservative and mutant residue. Each system was pre-equilibrated by pure MM energy minimization, followed by a 5 ns production MD run while the active site configuration was kept frozen. Then, the QM forces were activated, and 0.005 ns QM/MM trajectory was computed with frames being stored every 50 fs. The latest 50 frames underwent binding energy calculation, and the result was averaged. To perform QM/MM-PBSA, we adopted the MMPBSA.py script^118^ provided with the AmberTools package^119^. ATP with magnesium ion and catalytic and orientational water molecules were considered as a ligand, and the resting protein was considered as a receptor. All Poisson-Boltzmann and SASA/mbondi2 parameters were set to default^120^.

## Data availability

Raw sequencing reads for Hi-C and RNA-seq libraries, along with processed Hi-C maps in *mcool* format, loops in *bedpe* format, TPM-normalized RNA-seq genome-wide signal in *bigwig* format, and read count matrix, generated in this study, are available as a GEO SuperSeries under accession number GSE247397. token: gfutckuohnofrkv

## Code availability

The analysis code used to produce and analyze data has been made publicly available on GitHub: https://github.com/irzhegalova/dicty_cgp_loops

## Acknowledgements

We are grateful to Ilya Flyamer, Ed Banigan, Maxim Imakaev, Nezar Abdenur, and Leonid Mirny for productive discussions and help in the preliminary data analysis at the early stages of project development; to Nikolai Safronov for critical reading. We thank the Center for Precision Genome Editing and Genetic Technologies for Biomedicine, IGB RAS. This study was supported by the Russian Science Foundation (grant #21-64-00001 to S.V.R.) and the Russian Foundation for Basic Research (bioinformatic analysis; grant #20-34-90058 to I.V.Z.).

## Author contribution

S.V.R., A.A.Gav., and S.V.U. conceived the study. S.V.U. performed *D. discoideum* cultivation and induction of multicellular development. S.V.U. and P.A.V. prepared Hi-C libraries. M.D.L. performed RNA-seq. O.V.T. and A.A.Gal. performed Hi-C, RNA-seq and ATAC-Seq data processing and initial analysis, supervised by M.S.G. I.V.Z. analyzed Hi-C, RNA-seq, ATAC-seq and ChIP-seq data, supervised by M.S.G. I.A.P. performed logistic regression and analysis of Rabl configuration, supervised by E.E.K. A.V.L., and I.V.Z. annotated loops, supervised by S.V.R. I.A.P., I.V.Z., and S.V.U. annotated las-loops. A.A.Gal. performed polymer simulations. I.V.Z. annotated amino acid substitutions in *D. discoideum* cohesin ABC-like ATPse domains, supervised by S.V.U. E.S.B. performed molecular dynamics simulation of the cohesin subunits, supervised by D.N.I. S.V.U., I.V.Z., A.A.Gal., and E.S.B. wrote the initial manuscript draft and prepared figures. All authors discussed the results, commented on the draft and provided critical feedback throughout.

## EXTENDED FIGURE LEGENDS

**Extended Figure 1. a**, Size distribution of DpnII restriction fragments in *D. discoideum*, *S. cerevisiae*, *D. melanogaster*, and *H. sapiens* genomes. GC-content of the genome is shown above the boxplots. **b**, Clustering of Hi-C replicates. **c**, Whole-genome Hi-C maps at Dicty development stages. **d**, Eigenvector profile across Dicty chromosomes. **e**, Network of las-loops identified in vegetative cells (0h) and late aggregates (8h).

**Extended Figure 2. a**, Size distribution (upper panel) and proportion (bottom panel) of consecutive and stand-alone loops. **b**, Pileups of IS profiles (related to Fig. 2d). Data source: *S. cerevisiae* - Micro-C XL in S-phase cells^51^; *D. melanogaster* - Hi-C in spermatogonia^52^; *H. sapiens* - Hi-C in GM12878 lymphoblastoid cells^50^.

**Extended Figure 3. a,** Clustering of RNA-seq replicates. **b,** A heatmap of gene expression at Dicty development stages. One row represents one gene. Genes are ordered according to hierarchical clustering. **c,** UMAP of loop k-means clustering according to the coverage of genes with the same developmental trajectories. **d,** A scheme of the genes clustering into TGCs and grouping of loops by the TGC coverage.

**Extended Figure 4. a-d**, Pileups for Fig. 4d-f. **e**, Dependency between changes of insulation score (IS) and transcription in 2-kb bins in comparison between vegetative cells (0h) and late aggregates (8h). **f,** Averaged Hi-C maps centered at genes downregulated (left panel) and upregulated (middle panel) at 2h stage compared to 8h stage, and at genes ubiquitously expressed at both stages (right panel). **g,** Related to Fig. 4h: the same picture, but groups of loops are defined by 25^th^ percentiles (left) and 35^h^ percentiles (dashed lines) of transcription level around loop anchors to demonstrate the robustness of the analysis.

**Extended Figure 5. a,** CGP enrichment at loop anchors and inside loops. **b**, Size distribution of ELEs. **c,** Distributions of expression level of genes which contain and do not contain ELEs. *p*-value in the MWU. **d,** Representative examples of ELEs. H3K27ac, H3K4me1, ATAC-seq profiles and gene positions are shown. **e,** Averaged interaction between DGPs separated by different genomic distances.

**Extended Figure 6.** Averaged Hi-C maps centered at divergent gene pairs (DGPs) with different levels of transcription (genes transcribed at high and low levels are highlighted with dark gray and light gray, respectively).

**Extended Figure 7.** Preliminary sweep and local optimization of polymer simulation parameters. **a,** Quality metric for the preliminary parameters sweep. As Dicty loops are non-hierarchical, we used the median value of N loop strength from the experimental data, while keeping N+1 loop strength at around 1. Scheme of N and N+1 hierarchical loops (leftmost), loop strength in experimental Hi-C data (central left; blue zone - target values for the strength of N loops, red zone - target values for the strength of N+1 loops), model polymer used for preliminary parameter sweep (central right), loop strength distributions in the model polymer under different parameters (rightmost). Circles in different colors highlight loops in the model polymer used as a quality measurement, red arrows represent convergent gene pairs (CGPs). **b,** Transcription initiation rate vs extruder/RNA-polymerase speed ratio preliminary parameter sweep using strength of average N loop (left) and strength of average N+1 loop (right) as target values for optimal parameter windows selection. Optimal values falling into the inter-quartile range of Hi-C loop strength are designated with blue rectangles. **c,** Local parameter optimization using random genomic locus (chr4:750,000-1,000,000). Pearson correlation between experimental and simulated Hi-C maps is a quality metric for simulation performance. Transcription initiation rate and contact radius vs extruder lifetime in kb (left), extruder/RNA-polymerase speed ratio and transcription initiation rate vs extruder lifetime in kb (right). Optimal values are designated with blue rectangles.

**Extended Figure 8.** Distributions of correlations between simulated and experimental Hi-C data averaged over 244 partially overlapping 250-kb intervals covering the entire Dicty genome. **a,** Simulations performed at the extruder/polymerase speed ratio equal to 0.1, extruder lifetimes 10, 20 and 30 kb are tested. **b,** Simulations performed at the extruder lifetime 30 kb, extruder/polymerase speed ratios equal to 0.1, 1 and 10 are tested. **c,** Distribution of averaged correlations obtained at optimal simulation parameters: extruder/polymerase speed ratio = 0.1, extruder lifetime = 30 kb, transcription initiation rate = 0.05, contact radius = 1. Correlation has been calculated for contacts less or equal to 50 kb. **d,** An example of the genome locus whose structure is poorly predicted by simulations.

**Extended Figure 9.** Dicty cohesin substitutions. **a,** Tree of Life from iTOL, Dicty is colored in red. **b,** Phylogenetic tree of SMC3 and SMC1. SMC1 is highlighted in pink, SMC3 is highlighted in blue, Dicty is colored in red. **c,** Resulting values in molecular dynamics modelling, columns: Dicty substitution, gas-phase QM/MM energy difference change, polar solvation energy difference change computed by solving Poisson-Boltzmann equation, non-polar solvation energy change calculated in terms of SASA/mbondi2 model.

