## Supplementary figures and images for "Convergent gene pairs restrict chromatin looping in *Dictyostelium discoideum,* acting as directional barriers for extrusion"

### Extended Fig. 1

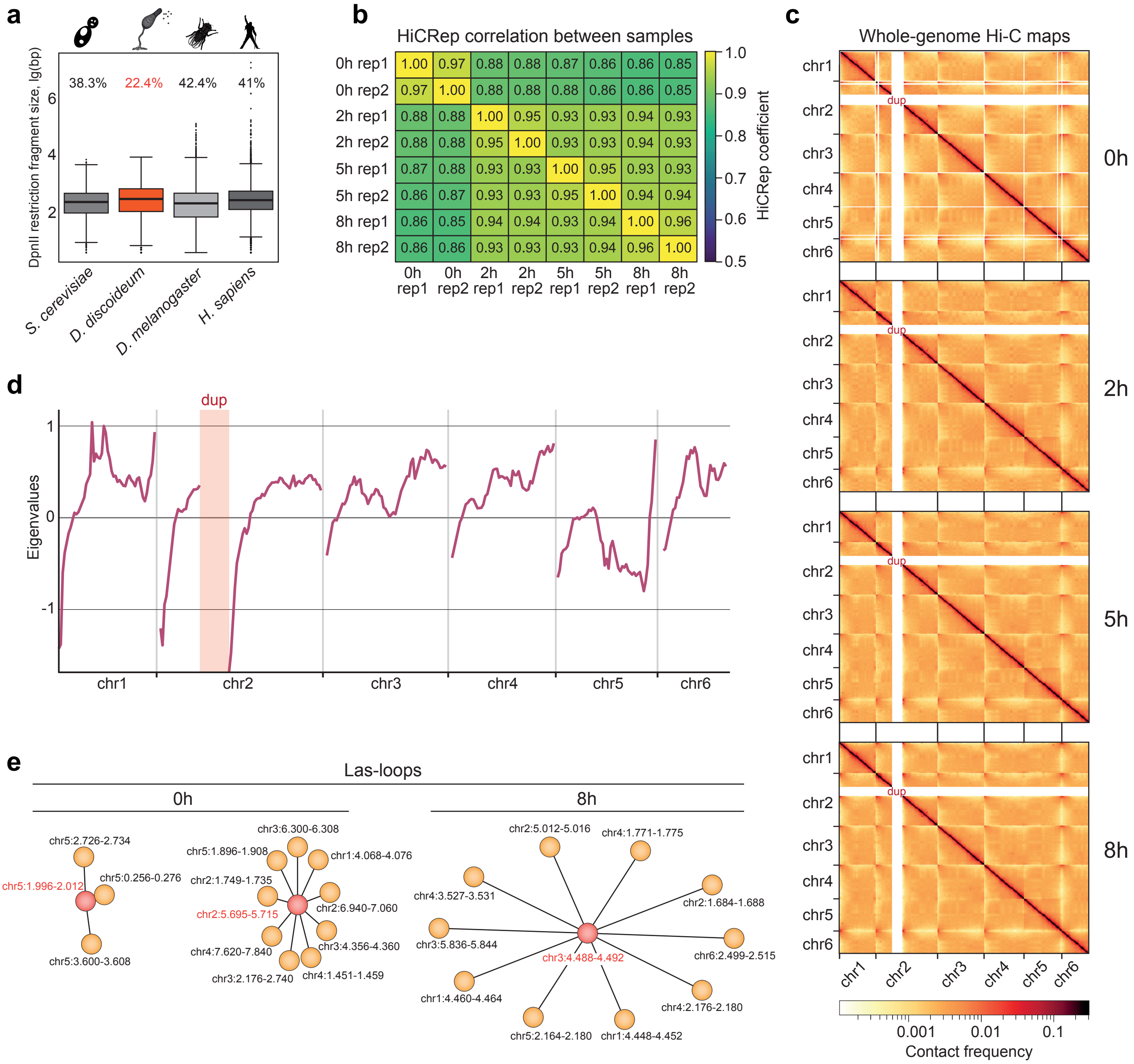

### Extended Fig. 2

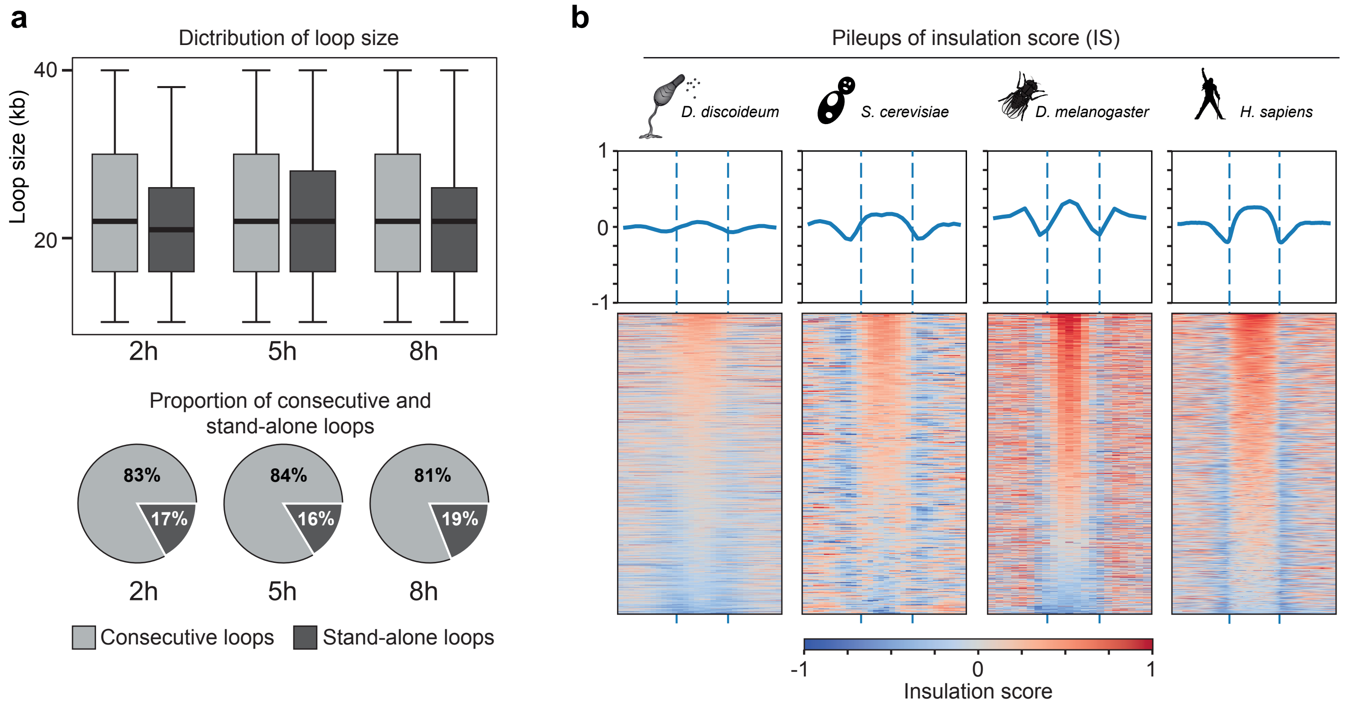

### Extended Fig. 3

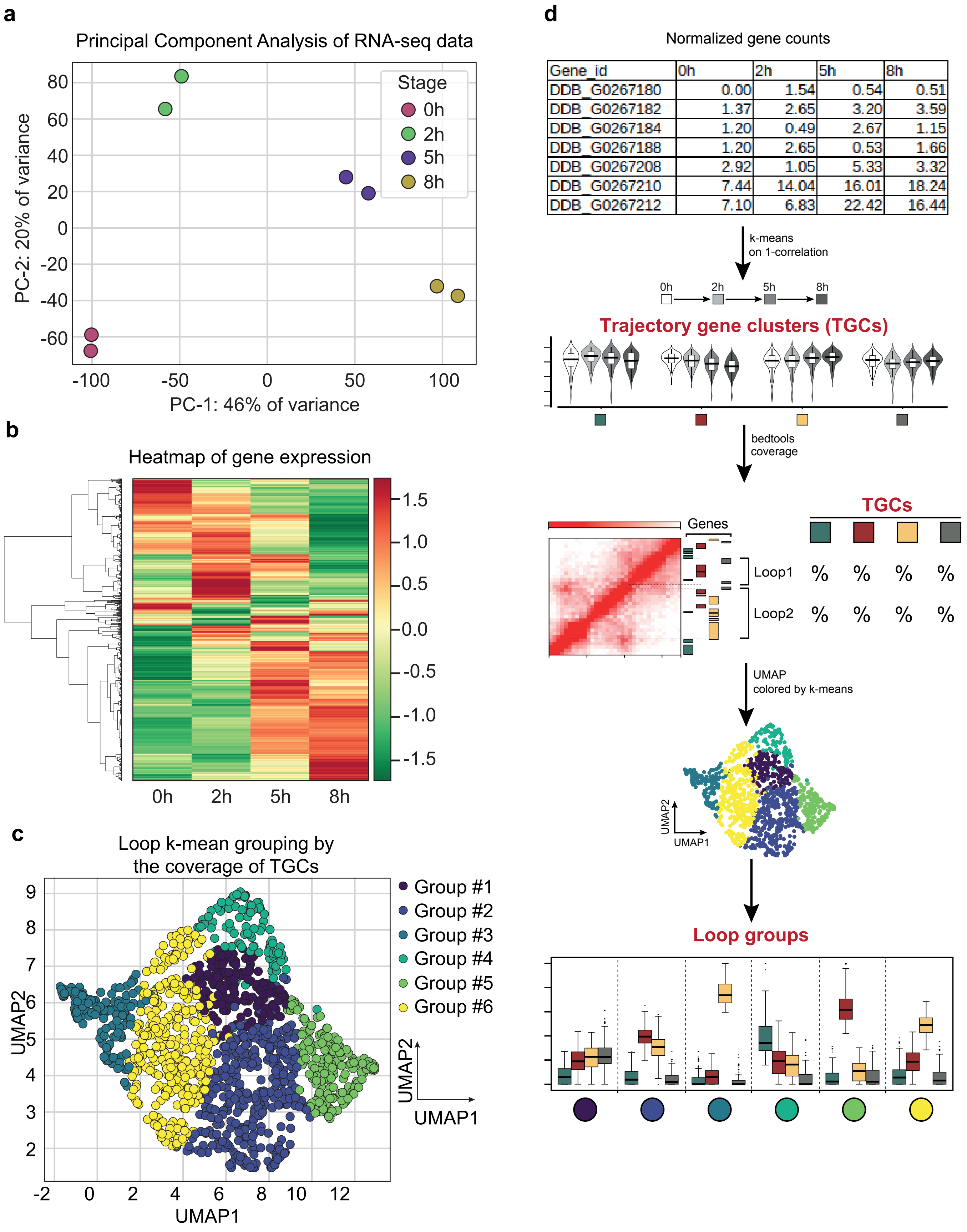

### Extended Fig. 4

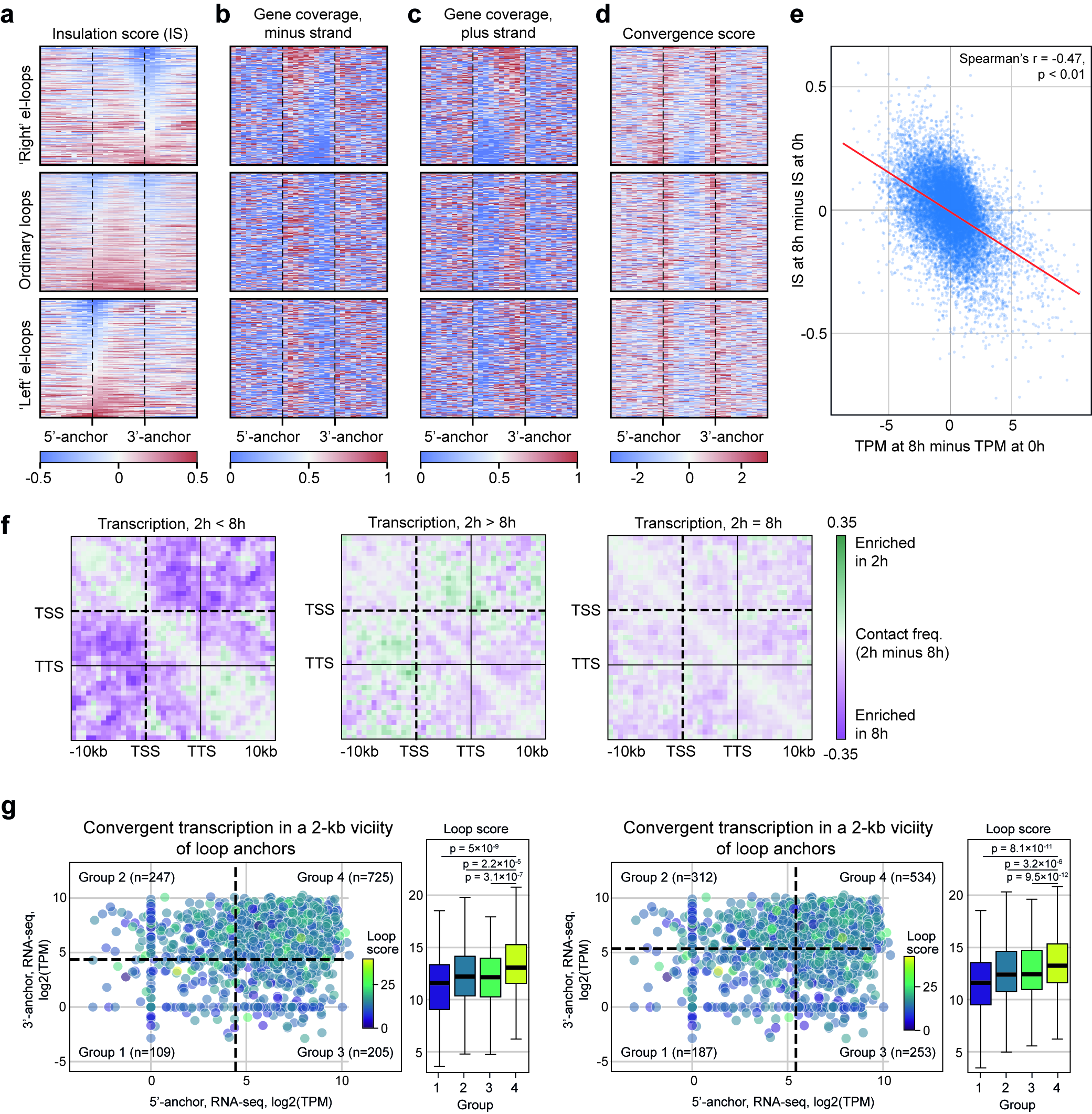

### Extended Fig. 5

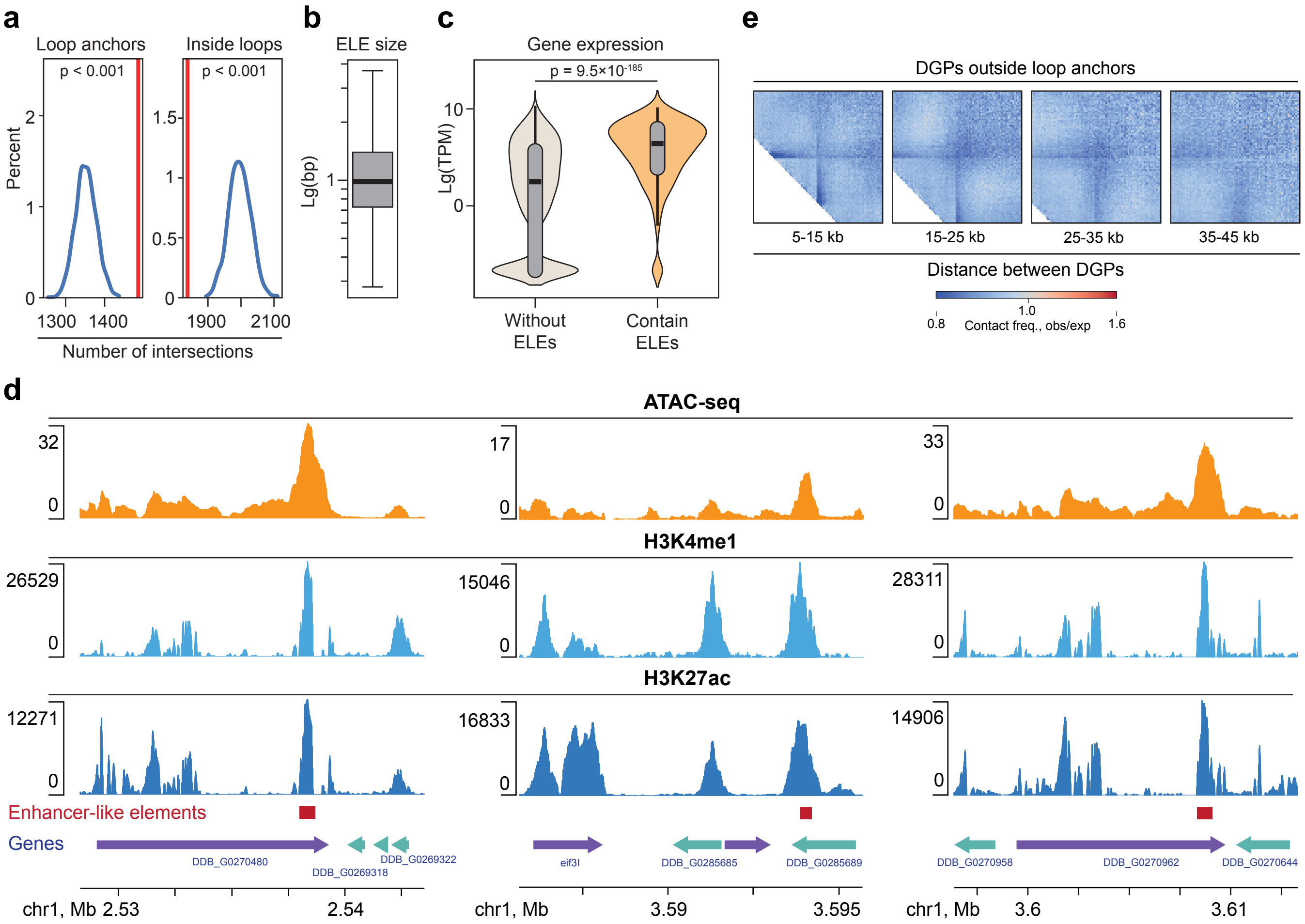

### Extended Fig. 6

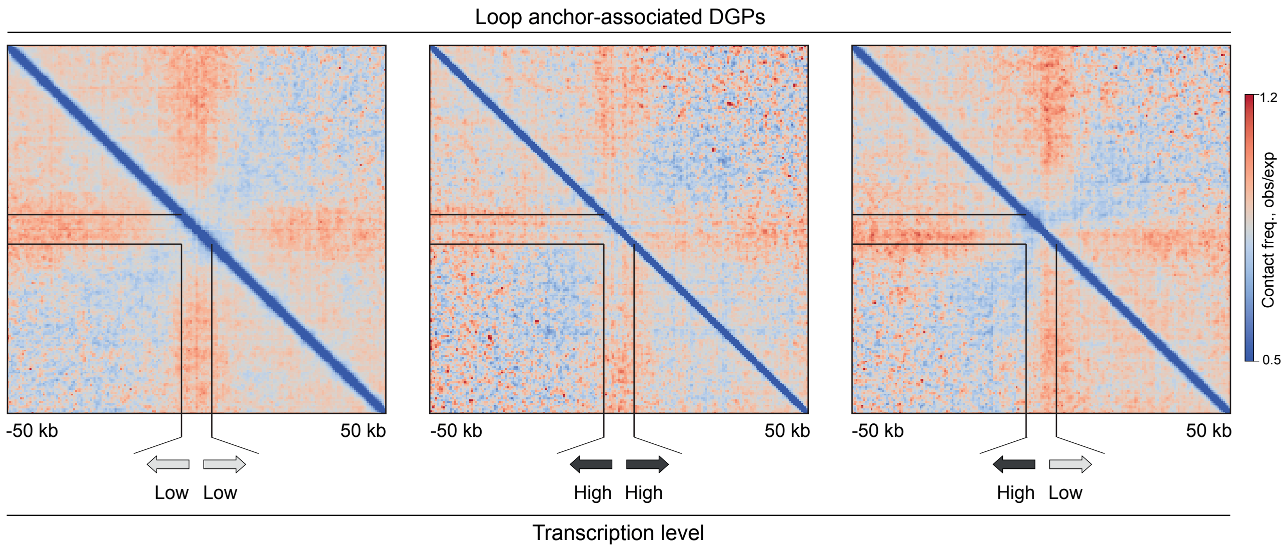

### Extended Fig. 7

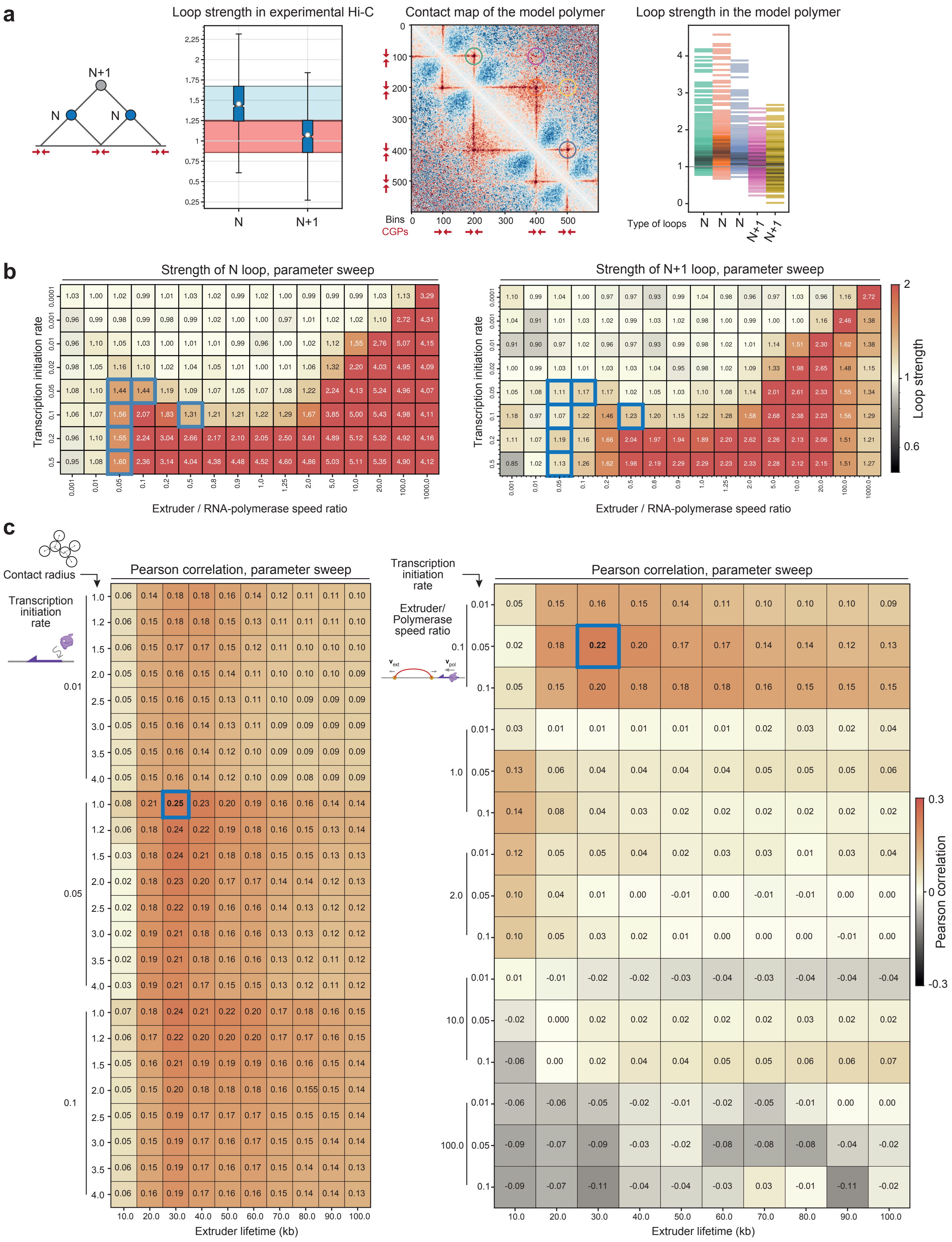

### Extended Fig. 8

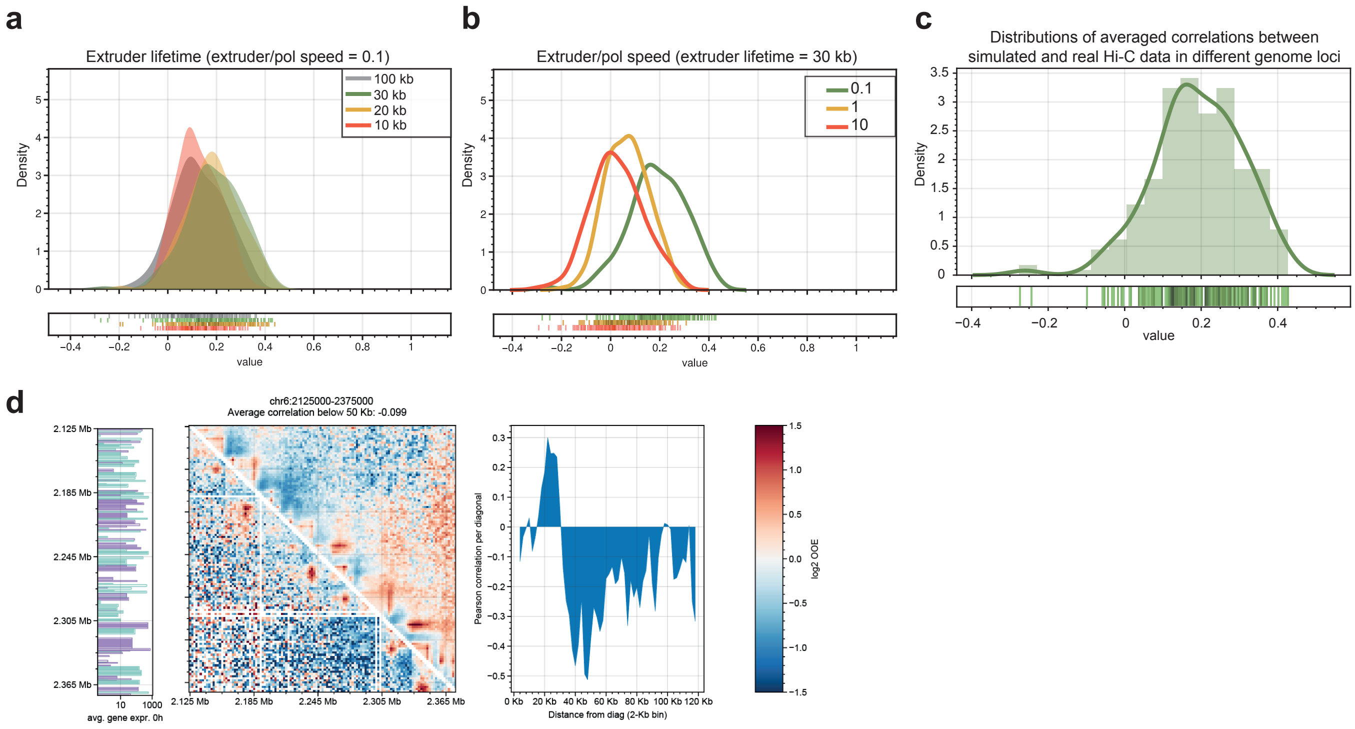

### Extended Fig. 9

a

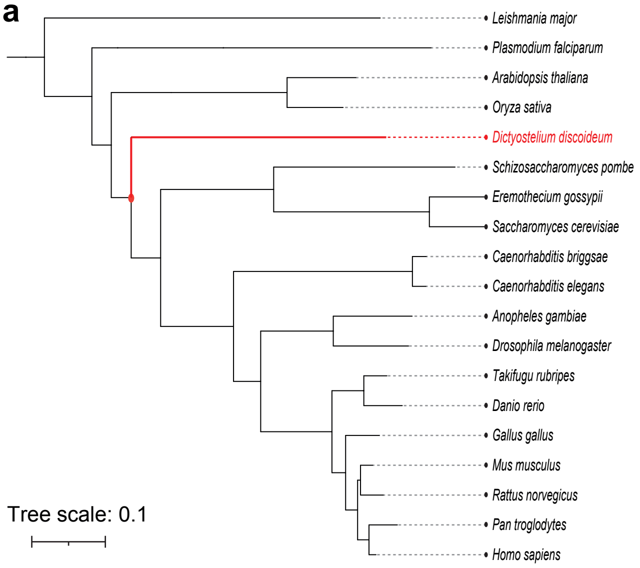


b

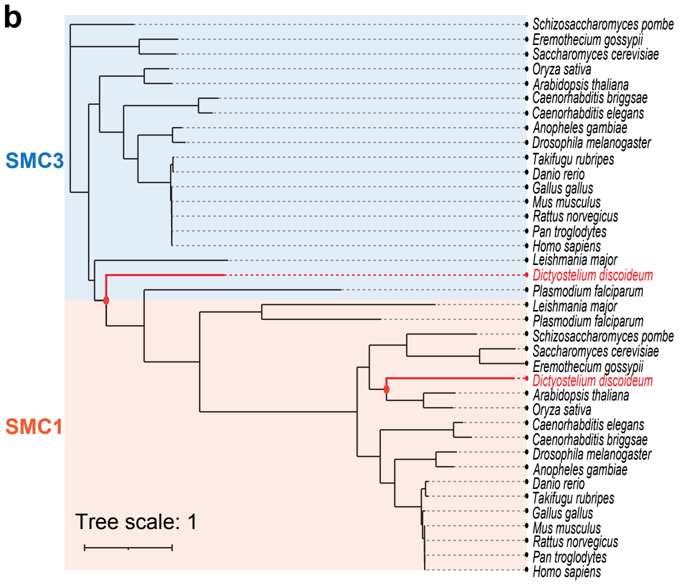


c

| Subsitution | ΔΔE(QM/MM) | ΔΔG(PB) | ΔΔG(SA) | ΔΔG(bind) |
|---|---|---|---|---|
| S37A | 1.1±0.2 | -0.2±0.1 | -0.1±0.0 | 0.8±0.2 |
| S35R | 1.6±0.4 | -0.1±0.2 | 0.4±0.2 | 1.9±0.4 |
| V1270I | 0.0±0.0 | -0.1±0.0 | 0.0±0.1 | -0.1±0.1 |
| L1278F | 0.0±0.1 | 0.1±0.2 | 0.0±0.2 | 0.1±0.2 |
| V1270I+L1278F | 0.0±0.3 | 0.1±0.1 | 0.0±0.1 | 0.1±0.3 |
