## Supplementary for "Convergent gene pairs restrict chromatin looping in *Dictyostelium discoideum,* acting as directional barriers for extrusion"

### SUPPLEMENTARY MATERIALS

**Supplementary Table 1.** Sequencing statistics of Hi-C libraries.

**Supplementary Table 2.** Las-loops identified in the vegetative (A) and late aggregation stages (B).

**Supplementary Table 3.** Loop coordinates in vegetative cells, migrating, and aggregating Dicty cells. The following parameters are listed: coordinates of the 2-kb genomic bin identified as the 5'-anchor (chrom1, start1, end1), coordinates of the 2-kb genomic bin identified as the 3'-anchor (chrom2, start2, end2), loop score (the same as the loop strength), loop type (elongated or regular), development stage.

**Supplementary Table 4.** Gene expression levels in vegetative, migrating, and aggregating Dicty cells, separately for replicates.

**Supplementary Table 5.** Loops containing genes with highly similar networks of gene ontology terms, as revealed by the best match average (BMA) analysis (BMA-loops). The following parameters are listed: loop coordinates (chrom, start, end), fold change over the random control, log10 of Mann-Whitney *p*-value, Kruskal-Wallis *p*-value; median BMA-value for all gene in loops, key GO terms.

**Supplementary Table 6.** Convergent (A) and divergent (B) gene pairs in the Dicty genome. The following parameters are listed for CGP: chrom, start, end of CGP; start and end of 5'-gene, start and end of 3'-gene, gene IDs, expression levels in TPM, length of genes, anchor group based on loop anchors intersection with CGP intergenic, log10 of intergenic length and sum gene length, group based on length both of the intergenic and gene length. The following parameters are listed for DGP: chrom, start, end of DGP; start and end of 5'-gene, start and end of 3'-gene, gene IDs, expression levels in TPM, intergenic length, expression group based on expression levels of both genes, anchor group based on loop anchors intersection with DGP intergenic,

**Supplementary Table 7.** Enhancer-like elements (ELEs) in the Dicty genome. Intergenic ELEs are designated by empty cell in the Gene ID column. The following parameters are listed: ELE coordinates (chrom, start, end), ID of the encompassing gene.

**Supplementary Table 8.** Dicty cohesin subunits: gene name, Dicty gene IDs, Uniprot ID, protein existence evidence from UniProt, and TPM-normalized transcription levels at four development stages. Median expression levels for all Dicty genes and beta-tubulin is also shown for comparison purpose.

**Supplementary Table 9.** Assessment of the RNA-polymerase density on Dicty chromatin and expected average initiation rate at TSS. Number of polymerases per kb of a gene is assumed based on yeast studies[1].

**Supplementary Table 10.** Preliminary optimization of parameters. The following parameters are listed: extruder processivity, extruder/polymerase speed ratio, transcription initiation rate, contact radius and loop strength.

**Supplementary Table 11.** Local optimization of parameters. The following parameters are listed: extruder processivity, extruder/polymerase speed ratio, transcription initiation rate, contact radius, average Pearson and Spearman correlations.

**Supplementary Table 12.** Whole-genome parameter optimization. The following parameters are listed: extruder processivity, extruder/polymerase speed ratio, transcription initiation rate, contact radius, region coordinates, average Pearson and Spearman correlations.

### Supplementary Methods

#### Polymer simulations of 3D genome

*Bead-Spring Polymer Model*

We used the classical bead-spring model using `OpenMM` (v7.7)[2] and the `polychrom` toolkit (v0.1.0) (https://github.com/open2c/polychrom), as described in previous studies[3-5] using publicly available code as a basis (https://github.com/mirnylab/moving-barriers-paper/tree/main). In brief, each DNA monomer is represented as a bead connected to neighboring genomic regions by a spring. The polymer chain of connected beads forms a linear chromosome. Polymer chain is placed into periodic boundary conditions.

A genomic region of 250 bp was selected as a single monomer, because: (1) it roughly corresponds to the DNA wrapped around a nucleosome, (2) it is smaller than the best resolution of Hi-C maps for Dicty (2 Kb), and (3) it is smaller than the average sizes of genes and intergenic regions (~1540 bp and ~670 bp, respectively), allowing most genes and intergenic regions to be represented by several monomers. Additionally, a typical Dicty chromosome (e.g., chr5, ~5.1 Mb) can be represented as ~20,000 monomers of this size, making simulations feasible and relatively fast on standard GPUs.

The polymer was initialized as a random walk on a cubic lattice using polychrom's `grow_cubic` function with a density of 0.1. The bond distance was set to 1.0 with a 0.1 wiggle distance. The angle force for the polymer chain was set to 1.5, and repulsion between monomers was set to the "trunc" parameter at 1.5, allowing chain crossing.

Extruders were simulated as additional bonds between looped monomers, with a bond length of 0.5 and a 0.2 wiggle distance. The extruders did not alter the polymer's other properties. Polymerases were not explicitly simulated in the bead-spring model.

The polymer dynamics were simulated with a variable Langevin integrator, with an error tolerance set to 0.01. The interaction of the polymer with the surrounding solvent was simulated with a collision rate of 0.03, following previous works[3].

*Simulations of Extruder Movement and Interaction with Polymerase*

Before running polymer simulations, we sampled trajectories of extruder and polymerase movement on a linear DNA lattice, where the grid size corresponded to the bead size in 3D simulations. To simplify calculations, we simulated lattice walkers on a single DNA region with a small system size (300 monomers or 75 Kb for the "Four Convergent Gene Pairs Model" and 1000 monomers or 250 Kb for the "Genomic Locus Simulations") and repeated it 50 times to model the whole chromosome.

Each extruder had two sides (left and right), each occupying one grid element on the lattice (Suppl. Fig. 1a, top). Extruders loaded on a single grid element and moved in opposite directions unless encountering an obstacle (another extruder) or being pushed by a polymerase. Extruders could spontaneously detach with a probability defined by the lifetime parameter, immediately re-attaching to a random lattice position. The pool of extruders was limited and defined by the separation parameter. We assumed uniform extruder loading, although a bias towards open chromatin regions is possible[6].

Each polymerase occupied one grid element on the lattice, loading at the gene start and progressing towards the transcription end site (Suppl. Fig. 1a, top). Polymerases did not stall, and their pool was not limited. The number of polymerases on the lattice is defined by the polymerase loading rate at transcription starts (probability of polymerase loading at a given position at each simulation step). Polymerases in our simulations did not pause at transcription start sites (TSSs), terminated strictly at termination sites, and detached immediately thereafter (although promoter-proximal pausing of polymerase is likely in Dicty[7]).

The lattice simulation steps corresponded to single movements of extruder sides (when $vleg \geq vpol$) or polymerase movements (when $vleg < vpol$), depending on their relative speeds. When $vleg \geq vpol$, the speed of polymerase is defined as a number from 0 to 1 that corresponds to the probability of stepping of polymerase at each step of extruder. When $vleg < vpol$, the speed of extruder leg is defined as a number from 0 to 1 that corresponds to the probability of stepping of extrider side at each step of polymerase.

Extruders encountering polymerases in head-to-head collisions were stopped and pushed back by the polymerase in the direction of the polymerase movement (Suppl. Fig. 1a, central top). In head-to-tail collisions, extruders moved forward with the polymerase speed (this scenario is only possible when vleg>vpol). Extruders overtaken by polymerases in tail-to-head collisions were pushed by the polymerase (this scenario is only possible when vleg<vpol , Suppl. Fig. 1a, central bottom).

In our simulations: (1) extruder lifetime was unaffected by collisions with polymerases, and (2) extruders could not bypass polymerases unless they detached at the gene end. We explored these scenarios by introducing and adjusting corresponding parameters but found no substantial improvement in model fitting (data not shown).

*Simulations of 3D Dynamics of Polymer Model*

We converted 1D lattice walkers into a dynamic 3D polymer model. Each extruder corresponded to an additional bond in the polymer chain, with bond locations changing with each 1D step. We ran 200 polymer dynamics steps per 1D step and stored the 3D conformation at each 1D step. In total, we ran 11,000 steps of 1D dynamics for each simulation in "Four Convergent Gene Pairs Model" and "Genomic Locus Simulations".

*In silico Hi-C Map Generation*

For each simulation run, we took all generated 3D conformations, identified bead pairs within the contact radius, and counted them using the `monomerResolutionContactMapSubchains` function of `polychrom`. The contact radius was one of the parameters optimized in our simulations (see below).

*In silico* Hi-C maps were treated as regular Hi-C maps. We calculated observed-over-expected maps after normalizing by per-diagonal expected values using the cooltools function `observed_over_expected` (`lib.numutils`)[8] for genomic region simulations or custom Python code for toy simulations. For comparison to experimental Hi-C, we coarse-grained *in silico* maps to 2-kb resolution using `cooltools.lib.numutils.coarsen`. Whole-genome simulation contacts were stored as `cool`-files with `cooler.create_cooler`[9] and visualized in the Resgen Higlass-based 3D genome browser[10].

We assumed that *in silico* Hi-C does not have any unmappable regions (e.g., telomeric regions or repeats at TSSs known for Dicty[7]), and does not have any sequencing biases[11], with all observed polymer contacts resulting in *in silico* Hi-C map contacts without sampling. Although we tried to simulate sampling of contacts due to experimental Hi-C biases, it did not improve the quality of Hi-C prediction based on simulations (data not shown).

*Initial Parameter Range Choice*

To parameterize the rules for this minimal model, we considered several observations. RNA polymerase speed for Dicty has been assessed at 1.3 kb/min[12], similar to *Drosophila* (~1 kb/min)[13] and mammals (1.25-3.5 kb/min)[14], but faster than yeast RNA polymerase (~0.8 kb/min)[15]. Using yeast data on RNA polymerase II density (0.078 mol/kb in genes)[1], we estimated approximately one RNA polymerase molecule per 17.4 kb of the Dicty genome, with an expected average gene initiation rate of 0.115 (see “Methods: Estimating number of RNA polymerase per TSS” and Supplementary Table 9). Despite reports that cohesin can bypass polymerase[16], we assumed polymerase acts as an impermeable barrier to extruders in Dicty simulations in both heads-on and co-directional encounters[4]. We also assumed that extruder processivity was close to the linear separation between extruders, similar to previous assessments in mammals[17,18].

We made weak assumptions about extruder speed. Studies suggest condensin movement can range from 60 bp/s[19] to 1.5 kb/s[20], with some simulations using even larger speed ratios[3].

*Estimating number of RNA polymerase per TSS*

The RNA polymerase rate of synthesis for Dicty is 1.3 kb/min[12]. With an average size of the *Dictyostelium* gene of 1.5 kb, this suggests that one polymerase travels the gene in ~69 seconds, slightly more than a minute. The density of polymerases was not estimated in Dicty, but suggesting the similarity of genome sizes, gene density, and polymerase speed, the closest guess could be the polymerase density from yeast *S. cerevisiae* (0.078 mol/kb)[1]. This estimate suggests that, given the gene coverage of 73.5% out of the 33.95 Mb genome, there are ~2000 polymerases working in the Dicty genome (or one polymerase per 17.4 kb of the genome). With 0.681 TSSs per kb of transcribed genome, this results in an average initiation rate at TSS of 0.115, see Supplementary Table 9.

*Four Convergent Gene Pairs Model*

We used these estimates for parameter optimization in toy simulations of convergent genes. The simulated system was 300 beads or 75 Kb, repeated 50 times to produce a 3.75 Mb chromosome.

Genes were in convergent orientations with TSSs at positions 93, 107, 193, and 207. Each gene was 6 beads long (1,500 bp, close to average in Dicty), with intergenic regions of 3 beads (750 bp, close to 670 bp average in Dicty). The spacing between CGPs was 100 beads, or 25 kb, which is close to the estimated median loop size in Dicty.

We estimated the goodness of fit of simulated lop with Hi-C ones. The procedure was defined as follows.

(1) We first calculated in silico Hi-C matrices. We used “extended” regime where we looked not only at the contacts within 300-bead repeated regions, but also at the interactions between them. This allowed us to compute also the strength of N+1 and N+2 loops between CGPs (and the simulated region, thus, corresponded to 150 kb, see Extended Figure 7a, central right).

(2) We then calculated the observed over expected (o.o.e.) maps and the strength of loops (N, N+1 and N+2) between the CGP positions (where the position of CGP was defined as the bead between two TESs of convergent genes). The strength was estimated as the median of the o.o.e. contacts in

30-bead window (7.5 kb) around the CGP pair. Loop strength was compared to the loop strength calculated from experimental Hi-C maps between N and N+1 loops (Extended Figure 7a, central left).

(3) Mean N loop strength in Dicty Hi-C was ~1.47 and mean N+1 loop strength was ~1.07. We, thus, selected, the empirical criterium of goodness of fit in four CGP model: (i) strength of each N loop in the *in silico* map should be no less than 1.27, and (ii) strength of each N+1 and N+2 loop should be no more than 1.27, where the number 1.27 is the midpoint between mean N and N+1 loop strength.

We varied extruder leg / polymerase speed ratio, transcription initiation rate (Extended Figure 7b), as well as extruder lifetime and contact radius for four CGP model (Suppl. Figure 1b):

extruder leg / polymerase speed ratio: {0.0001, 0.001, 0.01, 0.05, 0.1, 0.2, 0.5, 0.8, 0.9, 1.0, 1.25, 2.0, 5, 10, 20, 100, 1000, 10000}

transcription initiation rate: {0.0001, 0.001, 0.01, 0.02, 0.05, 0.1, 0.2, 0.5}

extruder lifetime: {20, 30, 40, 50, 60, 70, 80, 90, 100}

contact radius: {1.0, 1.2, 1.5, 2.0, 2.5, 3.0, 3.5, 4.0, 4.5, 5.0, 5.5, 6.0, 6.5}

We tried all possible combinations of these parameters and present two tables with marked optimal ranges (Extended Figure 7b and Suppl. Figure 1b).

While the broad range of extruder lifetimes from 40 to 100 and multiple contact radii from 1.0 to 6.0 produced suitable results, only one area of speed ratio speed ~ 0.1 was optimal (Extended Figure 7b, blue). Transcription initiation rate for the suitable ranges was from 0.05 to 0.5. We used this parameter sweep as a starting point for investigation of parameters for simulations of realistic genomic locus (see below).

*Genomic Locus Simulations*

We then noted that toy simulations between four CGPs do not represent all the variability of the gene and intergenic region sizes, distributions in the positioning and orientation of the genes. Thus, we extended our simulations to two random 250-kb regions of Dicty genome, #1: chr4:750,000-1,000,000 and #2: chr5:2,125,000-2,375,000, and devised simulations that simulate polymerase and cohesin movement at all genes in these regions, proportional to the RNA-Seq-measured transcriptional activity.

Using *bioframe*[21], we overlapped the selected genomic regions with non-overlapping genes, and selected the genes that are fully contained within the studies regions. We then converted the positions of these genes into 1D grid lattice with 250-bp grid element size, and assigned the relative polymerase initiation rate to each TSS proportional to the mean-normalized RNA-Seq signal on these genes. Note that relative polymerase initiation rate reflects relative to the average in the genome, while average initiation rate in simulations is varied separately by corresponding parameter (see above).

We then run polymer simulations for the corresponding genomic regions, and constructed *in silico* Hi-C maps for them, while sweeping extruder transcription initiation rate and contact radius (Extended Figure 7c left), as well as extruder lifetime and leg / polymerase speed ratio (Extended Figure 7c right).

For each simulation, we coarse-grained maps to 2 kb, calculated observed over expected (see "Methods: In silico Hi-C Map Generation"), compared each diagonal of *in silico* map to experimental Hi-C map by taking Pearson correlation (see Figure 7b "Correlation" panels), and then took average across distances from 6 Kb to 50 kb. Resulting maps of average by-diagonal Pearson correlation coefficients are shown in (Extended Figure 7c left, right).

We varied extruder leg / polymerase speed ratio, transcription initiation rate (Extended Figure 7c left), as well as extruder lifetime and contact radius for *in silico* Hi-C (Extended Figure 7c right):

extruder leg / polymerase speed ratio: {0.1, 1, 2, 10, 100}
transcription initiation rate: {0.01, 0.05, 0.1}

extruder lifetime: {10, 20, 30, 40, 50, 60, 70, 80, 90, 100

contact radius: {1.0, 1.2, 1.5, 2.0, 2.5, 3.0, 3.5, 4.0}

We tried all possible combinations of these parameters and present tables with marked optimal correlations (Extended Figure 7c): transcription initiation rate 0.05, leg / polymerase speed ratio 0.1, cutoff 1.0, and lifetime 30 (see Suppl. Fig. 1c for visualization of the result).

However, when we repeated the same procedure for other loci of the genome, we found that the optimal parameters were slightly different for other loci (data not shown). Thus, the parameters optimized for a single region are not necessarily the best for each genomic region. This might be due to (1) potentially imprecise and/or biased assessment of RNA polymerase presence on the genes by RNA-Seq, (2) biological factors that might vary across the genome and impact loop extrusion or polymerase properties, but not accounted for in our model (such as targeted loading, density of targeted loading sites, variable extrusion seed across the genome).

We thus divised whole-genome simulations of all 250-Kb regions in the genome, to find the globally optimal parameters of loop extrusion with polymerase as the moving barrier.

*Whole-Genome Simulations*

For whole-genome simulations, we split the genome in 250-kb windows that tile the whole genome and overlap by their half (thus, each genomic region participated in simulations twice). We excluded low-mappability regions (>20% of unmapped bins in Hi-C for Dicty's vegetative stage) and regions without genes, resulting in 244 genomic regions.

We then run polymer simulations for each of these regions, while sweeping the parameters:

extruder lifetime: {10, 20, 30, 100}

leg / polymerase speed ratio: {0.1, 1, 10}

Due to heavy computations needed for these simulations, we set the contact radius to 2.0, and varied lifetime while fixing the speed ratio to 0.1 and varied speed ratio while fixing lifetime to 30. We also randomly sampled the regions for simulations with other parameters and combinations (from the list in section "Methods: Genomic Locus Simulations"), but have not found any that will be better by comparing medians of correlation distributions (data not shown). We found that changes in either lifetime or speed ratio from optimal (lifetime 30, speed ratio 0.1) resulted in worse predictions by medians (Extended Figure 8 a, b).

Importantly, these globally optimal parameters resulted in the simulated map of the region from "Methods: Genomic Locus Simulations" that was visually similar to Hi-C map, and subjectively more so, despite the formally lower average correlation (Suppl. Fig. 1d, comp. to Suppl. Fig. 1c). Moreover, when we took one of the worst predicted region for the global optimal parameters, we found that visual similarity between *in silico* and experimental Hi-C maps was distinguishable in both cases (Extended Figure 8d). In summary, the globally optimal parameters were effective for whole-genome simulations, highlighting the challenge of finding a single parameter set suitable for all genomic regions.

Finally, leg speed to polymerase ratio of 0.1 corresponds to 21.6 bp/sec speed of extruder leg (assuming speed of PolII is 1.3 kb/sec[12]), and, thus, to ~40 bp/sec speed for the two-sided extruder.

#### Supplementary Figures

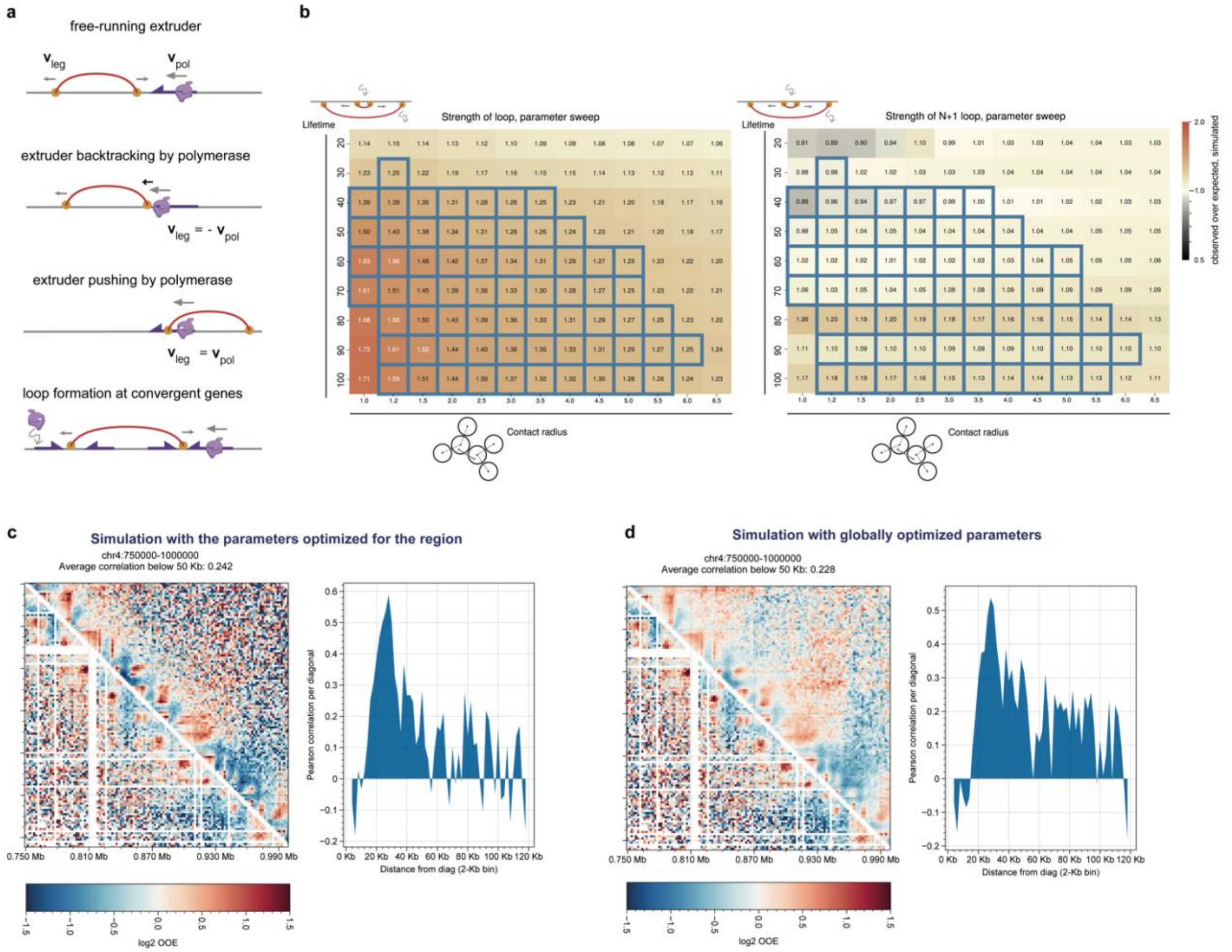


**Supplementary Figure 1. a**, Different rules for 1D extrusion interplay with polymerase movement. **b**, Lifetime vs contact radius cutoff for preliminary parameter sweep using strength of average N loop (left) and strength of average N+1 loop (right) as target values for optimal parameter windows selection. Optimal values falling into the interquartile range of Hi-C loop strength are designated with blue rectangles. **c**, Representative example of simulated Hi-C data (upper map) in comparison with experimental Hi-C data (bottom map) for a locus simulated with optimal parameters in Extended Figure 7c. Distributions of the Pearson correlation coefficient averaged over different genomic distances are shown to the right of the maps. **d**, Same locus as in Suppl. Fig. 1c, but simulated with globally optimized parameters.
